# Genomic variation across a clinical *Cryptococcus* population linked to disease outcome

**DOI:** 10.1101/2021.11.22.469645

**Authors:** Poppy Sephton-Clark, Jennifer L. Tenor, Dena L. Toffaletti, Nancy Meyers, Charles Giamberardino, Síle F. Molloy, Julia R. Palmucci, Adrienne Chan, Tarsizio Chikaonda, Robert Heyderman, Mina Hosseinipour, Newton Kalata, Cecilia Kanyama, Christopher Kukacha, Duncan Lupiya, Henry C. Mwandumba, Thomas Harrison, Tihana Bicanic, John R. Perfect, Christina A. Cuomo

**Affiliations:** Infectious Disease and Microbiome Program, Broad Institute of MIT and Harvard, Cambridge MA 02142 USA; Division of Infectious Diseases, Department of Medicine, Duke University School of Medicine, Durham, NC 27710, USA; Centre for Global Health, Institute of Infection and Immunity, St George’s University of London, London, United Kingdom; Clinical Academic Group in Infection, St George’s University Hospital, London, United Kingdom; Sunnybrook Health Sciences Centre, Toronto, Canada; Division of Infection and Immunity, UCL, London, United Kingdom; UNC Project Malawi, University of North Carolina, USA; Malawi-Liverpool-Wellcome Trust Clinical Research Programme, Blantyre, Malawi; Tisungane Clinic, Zomba Central Hospital, Zomba, Malawi

## Abstract

*Cryptococcus neoformans* is the causative agent of cryptococcosis, a disease with poor patient outcomes, accounting for approximately 180,000 deaths each year. Patient outcomes may be impacted by the underlying genetics of the infecting isolate; however, our current understanding of how genetic diversity contributes to clinical outcomes is limited. Here, we leverage clinical, in vitro growth and genomic data for 284 *C. neoformans* isolates to identify clinically relevant pathogen variants within a population of clinical isolates from patients with HIV-associated cryptococcosis in Malawi. Through a genome-wide association study (GWAS) approach, we identify variants associated with fungal burden and growth rate. We also find both small and large-scale variation, including aneuploidy, associated with alternate growth phenotypes, which may impact the course of infection. Genes impacted by these variants are involved in transcriptional regulation, signal transduction, glycosylation, sugar transport, and glycolysis. We show that growth within the CNS is reliant upon glycolysis in an animal model, and likely impacts patient mortality, as CNS yeast burden likely modulates patient outcome. Additionally, we find genes with roles in sugar transport are enriched in regions under selection in specific lineages of this clinical population. Further, we demonstrate that genomic variants in two genes identified by GWAS impact virulence in animal models. Our approach identifies links between genetic variation in *C. neoformans* and clinically relevant phenotypes and animal model pathogenesis; shedding light on specific survival mechanisms within the CNS and identifying pathways involved in yeast persistence.

**Importance:** Infection outcomes for cryptococcosis, most commonly caused by *C. neoformans*, are influenced by host immune responses, as well as host and pathogen genetics. Infecting yeast isolates are genetically diverse; however, we lack a deep understanding of how this diversity impacts patient outcomes. To better understand both clinical isolate diversity and how diversity contributes to infection outcome, we utilize a large collection of clinical *C. neoformans* samples, isolated from patients enrolled in a clinical trial across 3 hospitals in Malawi. By combining whole-genome sequence data, clinical data, and in vitro growth data, we utilize genome-wide association approaches to examine the genetic basis of virulence. Genes with significant associations display virulence attributes in both murine and rabbit models, demonstrating that our approach can identify potential links between genetic variants and patho-biologically significant phenotypes.

## Introduction

*Cryptococcus neoformans* is a pathogenic yeast that most commonly affects immunocompromised individuals, causing an estimated 180,000 deaths annually, with 75% of these occurring in sub-Saharan Africa. One of the leading causes of death in adults living with HIV/AIDS, cryptococcal infections are especially problematic in low-income countries where, despite a widespread roll-out of antiretroviral therapy, deaths due to opportunistic infections such as cryptococcal meningitis remain high (1). The infecting propagules of this pathogen generally enter human hosts via inhalation. From infections within the lung, *C. neoformans* may disseminate throughout the bloodstream and central nervous system of susceptible patients, causing life-threatening meningitis (2). In a sample of healthcare systems across low-income countries, the 1-year mortality rate for individuals who develop cryptococcal meningitis is estimated to be 70% for those in care (uncertainty interval 59–81%) (1). A better understanding of *C. neoformans* strain virulence and fitness within the host is necessary to improve patient outcomes and develop new treatment options.

Whilst the majority of cryptococcosis cases are caused by *Cryptococcus neoformans var grubii* (3), there are often high levels of genetic diversity within clinical populations of *C. neoformans* (4–7). Furthermore, isolates of the same multilocus sequence type (MLST) have been shown to cause infections that range in severity from mild to extreme (8). To examine how genetic variation contributes to virulence phenotypes, a recent study carried out logistic regression analysis with 38 clinical *C. neoformans* isolates of the same sequence type to identify single nucleotide polymorphisms (SNPs) associated with patient survival, clinical parameters including cytokine response, immune cell counts and infection clearance, as well as in vitro data on absolute yeast growth and macrophage interactions (9). This study identified 40 candidate genes based on these association parameters, 6 of which (out of 17 genes tested) were important for survival in a murine model of *C. neoformans* infection. In a larger cohort of 230 *C. neoformans* samples from patients in South Africa, isolate sequence type was associated with patient outcome, in vitro cerebrospinal fluid (CSF) survival, and phagocytosis response (10). Full scale genome-wide association studies (GWAS) have also examined how natural variation within a *C. neoformans* population differentiates clinical and environmental isolates, identifying loss-of-function variants present in clinical *C. neoformans* (VNB) populations that impact a transcription factor important for melanization, a well-studied virulence factor (11).

Furthermore, copy number variation, such as aneuploidy, has also been frequently identified within clinical populations of *C. neoformans*. Disomy of chromosome 1 is commonly reported for isolates exposed to azoles, and the higher copy number of two key genes, the *AFR1* transporter and the *ERG11* drug target, confer increased resistance to antifungals such as fluconazole (12–14). Chromosome duplication as a result of in vivo passage has also been noted in clinical isolates (15–17), and the emergence of aneuploidy in this setting has been proposed as a mechanism by which both *Cryptococcus* and *Candida* species might rapidly adapt to high-stress environments (18, 19). In *C. neoformans* aneuploidy is often transient and passage under non-selective conditions allows for reversion to euploidy (14, 17). In total, aneuploidy of chromosomes 1, 2, 4, 6, 8, 9, 10, 12, 13, and 14 have been reported in *C. neoformans* (14, 16, 17, 20–23). Despite appearing consistently throughout clinical populations, the impact of these other chromosomal aneuploidies is not yet well understood.

To better understand how genetic variation among *C. neoformans* isolates contributes to infection outcomes in patients, we carried out genome-wide association studies (GWAS) with 266 *C. neoformans* clinical isolates from the VNI lineage, selected to reduce the confounding effects of population structure between lineages. In addition to comparing selected clinical data, all isolates were also measured for in vitro growth under diverse conditions. Through our GWAS approach, we identify two proteins associated with fungal burden in patients and demonstrate a connection to virulence in animal models. Additionally, we show that yeast survival within the rabbit central nervous system (CNS) is dependent on a glycolytic gene identified by GWAS and corroborate findings that patient outcome is highly correlated with fungal burden in the CNS. Partial and full chromosomal duplications are commonly detected within this clinical population, yet these aneuploidies reduce *C. neoformans* fitness under in vitro growth conditions.

## Materials and Methods

### Sample Preparation and Sequencing

Clinical cryptococcal isolates derived from patient CSF subculture were procured through the Antifungal Combinations for Treatment of Cryptococcal Meningitis in Africa Trial (ACTA) (24); repeat cultures and duplicates were excluded. Collected strains were grown overnight in 10 mL of YPD at 30°C and 225 rpm. Genomic DNA was then extracted for sequencing with the MasterPure Yeast DNA Purification Kit, as described by Desjardins et al. (11). DNA was sheared to 250bp using a Covaris LE instrument and adapted for Illumina sequencing as described by Fisher et al. (25). Libraries were sequenced on a HiSeq X10, generating 150bp paired reads (minimum average coverage 100x).

### Data Processing and Variant Calling

To determine sample species, reads were first aligned to a composite pan-*Cryptococcus* genome, consisting of reference genomes for *Cryptococcus neoformans var. grubii* H99, *Cryptococcus neoformans var. neoformans* JEC21, and representative genomes for lineages VGI, VGII, VGIIIa, VGIIIb, VGIV, and VGV of *Cryptococcus gattii* (26–29). To identify variants for *C. neoformans* species, reads were aligned to the *Cryptococcus neoformans var grubii* H99 reference genome (GCA_000149245.3) (27) with BWA-MEM version 0.7.17 (30). GATK v4 variant calling was carried out as documented in our publicly available cloud-based pipeline (31) (https://github.com/broadinstitute/fungal-wdl/tree/master/gatk4). Post calling, variants were filtered on the following parameters: QD < 2.0, QUAL < 30.0, SOR > 3.0, FS > 60.0 (indels > 200), MQ < 40.0, GQ < 50, alternate allele percentage = 0.8, DP < 10. Variants were annotated with SNPeff, version 4.3t (32).

### Population Genomic Analysis

A maximum likelihood phylogeny was estimated using 72,258 segregating SNP sites present in one or more isolates, allowing ambiguity in a maximum of 10% of samples, with RAxML version 8.2.12 GTRCAT rapid bootstrapping (33), and visualized with ggtree (R 3.6.0) rooted to VNII isolates. Isolate lineage was identified based on phylogenetic comparison with previously typed isolates reported by Desjardins et al (11). To estimate linkage disequilibrium (LD) decay, vcftools version 0.1.16 was used to calculate LD for 1000bp windows, with a minimum minor allele frequency of 0.1, and the --hap-r2 option. Region deletions and duplications were identified using CNVnator v0.3 (significant instance e value < 0.01) (34). To identify regions under selection, composite likelihood ratio analysis was performed with PopGenome (R 3.5.0, PopGenome 2.7.5) per chromosome, by 5kb windows (35). The top 5% scoring regions (centromeric regions excluded) were tested for enrichment using a hypergeometric test with FDR correction. Large duplications and aneuploidies were visualized using funpipe (coverage analysis) version 0.1.0 (https://github.com/broadinstitute/funpipe).

### Genome-Wide Association Studies

Association analysis between clinical data, in vitro phenotypes, and variants was carried out using PLINK v1.08p formatted files and Gemma version 0.94.1 (36) (options: centered relatedness matrix gk 1, linear mixed model), as previously described (11). Variants were considered in two scenarios, one in which rare variants (present in < 5% of the population) were collapsed by gene, and another in which loss-of-function variants (SNPeff impact high) were considered independently. Significant variants were considered to have a test score < 1.00E-6.

### Clinical Data Analysis

De-identified clinical metadata detailing CSF fungal burden (CFU/mL), fungal clearance (EFA), patient mortality, and Glasgow coma score were provided by investigators, with these parameters determined as previously described (24). Correlation between clinical parameters was determined in R 3.6.0 with Pearson correlation coefficient, Spearman’s rank correlation coefficient, point-biserial correlation, or Phi coefficient. Survival curves were generated using Prism v9.1.0 and statistics were carried out in R 3.6.0.

### In vitro Phenotyping

Strains were grown at 30°C for two to five days. For each strain, a single colony was selected and added to 96 well microtiter plates containing 200 μL of YPD broth. Each 96 well plate contained six control strains (H99 and deletion mutants of *LAC1, MPK1, CNA1, RAS1*, and *HOG1*) and a YPD control. The 96 well plates were incubated for one to two days at 30°C. Colonies were pin replicated into 384 well microtiter plates containing 80 μL of YPD broth in each well. The 384 well plates were incubated for one to two days at 30°C. They were then pinned onto one well solid agar plates in duplicate using the BM5-BC Colony Processing Robot (S & P Robotics) in 1536 array format. In three separate biological replicates, isolates were grown at 30°C, 37°C, and 39°C on YPD agar. Isolates were also pinned onto YPD+10 μg/mL fluconazole and YPD+64 μg/mL fluconazole. Images were captured after approximately 24, 48, and 72 h. Colony size at 48h was determined using gitter (37), and used to assess growth.

### Gene Deletion Strains

Strains used for the animal studies and the primer sequences used are listed in Supplemental Table 1. KN99alpha (CM026) was used as the reference wild-type strain for deletions obtained from a genome-wide *Cryptococcus* deletion library (38). Multiple independent deletion strains were generated in wild-type *C. neoformans* strain H99 (*cnag_05608Δ, cnag_04102Δ, cnag_06033Δ*) for this study. Three DNA fragments were amplified by PCR: approximately 0.7-1 kb of 5’ flank sequence, the nourseothricin (NAT) drug selection cassette amplified from pAI3 (39), and 0.7-1 kb of 3’ flank sequence were prepared for each gene. The PCR products were gel extracted using the NucleoSpin Gel and PCR Clean-up Kit (Macherey-Nagel). Next, the PCR products were cloned into pUC19 using NEBuilder HiFi DNA Assembly, transformed into *Escherichia coli*, and positive plasmids were confirmed by PCR. For biolistic transformation, 2 μg of plasmid was transformed into *C. neoformans* strain H99 as previously described (40) with a slight modification that the yeast cells were recovered on YPD containing 0.5 M sorbitol and 0.5 M mannitol. The cells were allowed to recover for 2.5 h before transferring to the selective medium, YPD+100 μg/mL NAT. Positive transformants were confirmed by PCR.

### Capsule production

To evaluate the capsule size, capsule inducing medium (10% Sabouraud broth in 50 mM MOPS pH 7.3) was used as previously described (41). Five milliliters of capsule-inducing medium were inoculated with a single freshly streaked yeast colony and grown in an incubator shaker (225 rpm) for approximately 24 and 48 h. India ink was used as a counterstain at a 1:5 ratio (ink:cell suspension). Images were captured by microscopy (Zeis Axio Imager 1). Cell body and capsule size were measured in ImageJ V1.53a.

### Murine model of infection

*C. neoformans* strains were grown in YPD broth at 30°C in a shaking incubator (220 rpm) for 24 h, centrifuged, and washed twice in phosphate-buffered saline (PBS). Yeast cell counts were determined using a T4 cell counter (Nexcelom). For gene deletion strains, five male 22-24g CD-1 mice (Charles River Laboratories) were infected with approximately 5 × 10^4^ yeast cells per mouse via oropharyngeal aspiration while under isoflurane anesthesia. For clinical isolates, fifteen male 22-24g CD-1 mice (Charles River Laboratories) were infected with approximately 5 × 10^4^ yeast cells per mouse via oropharyngeal aspiration while under isoflurane anesthesia. Mice were monitored and weighed daily. Mice with a total body weight loss of ≥ 20% or that exhibited behavioral, neurological, or respiratory symptoms were sacrificed following IACUC guidelines, via CO2 asphyxiation. Kaplan-Meier survival plots and analysis (the log-rank test) were completed using Prism software v9.1.0.; GraphPad Software. A p-value of ≤ 0.05 was considered statistically significant. Outliers were excluded based on ROUT analysis performed using Prism software v9.1.0.; GraphPad Software.

### Rabbit model of infection

To assess the fitness and virulence of deletion strains in rabbit CSF, 3 New Zealand White male rabbits (2.3 – 4.7 kg) were inoculated intracisternally with 300 μL of approximately 1×10^8^ cells. Animals were sedated with ketamine and xylazine for inoculation and serial CSF cisternal taps. The rabbits were treated with hydrocortisone acetate (2.5 mg/kg) by intramuscular injections daily starting one day prior to yeast inoculation. Cisternal taps were performed on days 3, 7, and 10 followed by serial dilution of the CSF and enumeration of colonies. The time series fungal burden data were then assessed with estimated marginal means of linear trends using repeated measures analysis in R v3.6.1.

### Animal studies

Animal experiments were performed in compliance with the Animal Welfare Act, the *Guide for the Care and Use of Laboratory Animals* (42), and the Duke Institutional Animal Care and Use Committee (IACUC).

## Results

### The VNI lineage dominates clinical isolates and shows selection for sugar transporters

To examine variation within clinical populations, *C. neoformans* samples were isolated from HIV-infected patients as part of the ACTA trial and its human subject protocol (24), which evaluated the efficacy of fluconazole partnered with flucytosine, compared to amphotericin B combined with either fluconazole or flucytosine, as induction therapy for cryptococcal meningitis. Baseline (pre-antifungal exposure) isolates were collected from three hospitals in Malawi between 2013 and 2016. We performed whole-genome sequencing on 344 isolates and removed isolates identified as *Cryptococcus gattii* (45), hybrid AD *C. neoformans* (4), diploid (2), or with low coverage (9) from the analysis. To examine the population structure, a maximum likelihood phylogeny was built based on segregating SNP sites (Figure 1). Isolates can be clearly identified as VNI (266), VNII (9), and VNB (9), with VNI isolates split into VNIa (217) and VNIb (49) with 100% bootstrap support; these recently described sub-lineages show strong evidence of separation (11). Of the two mating type loci found in *Cryptococcus*, mating type □ predominated among these isolates, with only one VNB isolate (ACTA3525) possessing the alternate *MAT***a**. To assess recombination within the large VNI population, we calculated linkage disequilibrium (LD) decay and found levels of decay for lineage VNI (LD50 30kb) (Supplemental Figure 1) similar to those reported by Desjardins et al. (LD50 values for VNI, VNBI, and VNBII were less than 50kb) (11). There is increased decay in the VNI population as a whole when compared to individual VNIa and VNIb sub-groups (LD50 of 110kb for VNIb, LD50 not reached within 250kb for VNIa), suggesting that VNIa and VNIb isolates do not recombine exclusively within their groups.

**Figure 1.**
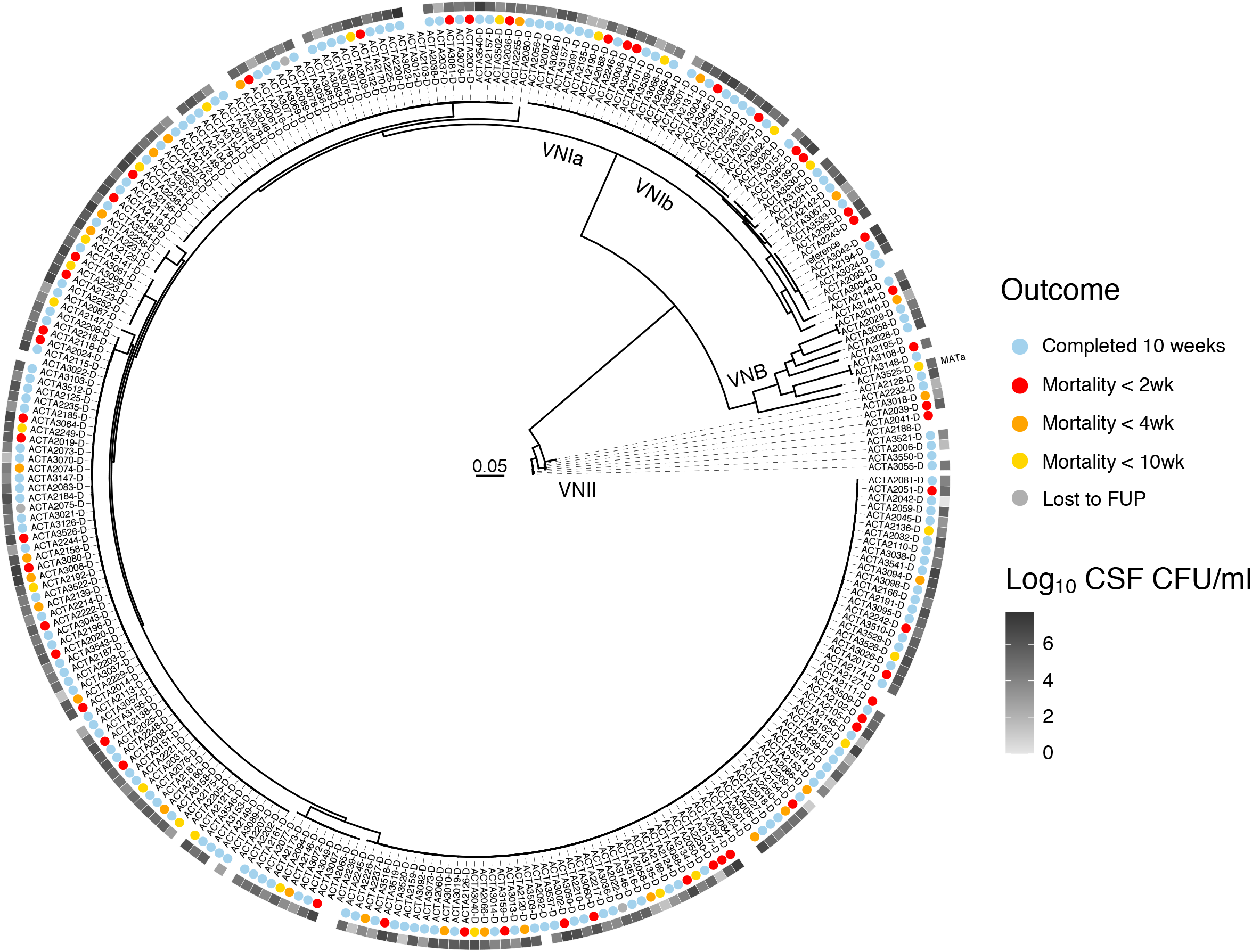
Maximum likelihood phylogeny of patient isolates, estimated from 72,258 segregating SNP sites, rooted to VNII. Isolates separate distinctly into VNI, VNB, and VNII, with all lineages having 100% bootstrap support. All isolates possess MAT⍰ except ACTA3523 (highlighted). Colored circles correspond to patient survival. Greyscale squares indicate patient fungal burden of cerebrospinal fluid prior to treatment, Log_10_(CFU/mL).

To identify genomic regions under positive selection, we performed composite likelihood ratio analysis (43). We found that regions with scores in the top 10% in more than one lineage (examining VNIa, VNIb, VNII, and VNB) include subtelomeric regions, centromeres, *ERG11*, and *AFR1* (Supplemental Figure 2). To examine if genes within regions showing high selection scores are associated with shared functions, we performed gene ontology (GO) enrichment analysis (Hypergeometric test, FDR correction) on regions with selection scores in the top 5% (excluding centromeres). For VNII isolates we found nucleotide excision repair enriched in these regions (corrected p-value = 6.77E-3). Sugar transport, including inositol transport, appeared robustly enriched in both VNIa and VNB lineages (corrected p-values = 1.07E-3 and 8.60E-3), supporting previous work that identified these functions as under selection (Supplemental Table 2) (11). To investigate the differences between functions under selection across lineages, we expanded our analysis of the VNII lineage to include genomic data from an additional 34 isolates for which we have whole genome sequencing data (11, 22). In this expanded VNII cohort, we found sugar transporters were enriched (corrected p-value = 8.5E-4) in regions with selection scores in the top 5% (excluding centromeres). As genes with roles in hexose transport have been identified to be enriched in the subtelomeric regions of *Cryptococcus* sp. chromosomes (44), we examined the locations of sugar transporters that appeared under selection in VNI, VNB, and VNII. In total, 65% of sugar transporters in regions under selection fall into subtelomeric regions; however, only two of these genes are predicted to function specifically as hexose transporters (Supplemental Table 2). Sugar transport and utilization has been identified as key to success in nutrient scarce environments such as the CNS (23, 45, 46), as important during interactions with amoebae, and for roles in virulence and resistance to external stress (47–50).

We also identified regions that were duplicated or deleted in these *C. neoformans* isolates via copy number variation analysis (Supplemental Table 3). An 8 kb region was found to be duplicated in 43 VNIa isolates, containing 4 genes including a sugar transporter (transporter classification database: 2.A.1.1, glycerol transport), a predicted non-coding RNA, a fungal specific transcription factor (Zn2Cys6, *SIP402*), and a short-chain dehydrogenase. A separate 34 kb region was found duplicated across 48 VNIa isolates encoding 11 genes including 3 dehydrogenase enzymes and 2 hydrolase enzymes. Duplicated regions unique to VNIb included an un-annotated 19kb region specific to 20 isolates and an 8kb region that encodes 2 hypothetical proteins duplicated across 21 VNIb isolates. Of the lineage-specific duplicated regions observed, all appeared to be specific to monophyletic groups within these lineages. Duplications of genes involved in resistance to azoles such as *ERG11* were found exclusively in VNII isolates; however, this finding did not correlate with an enhanced ability to grow in the presence of fluconazole at 64ug/mL (Supplemental Figure 3). While these duplicated regions are not directly associated with our tested phenotypes, the duplication of regions containing genes such as *ERG11* and sugar transporters may still contribute to phenotypic variation that is relevant to clinical outcomes through modulation of growth and virulence phenotypes when grown in alternate conditions.

### GWAS identifies multiple variants associated with fungal burden

We next used clinical data associated with these isolates to investigate the relationship between clinical factors, across lineages. We confirmed previous findings that mortality correlates strongly with high baseline fungal burden (log_10_ CSF CFU/mL taken at diagnosis of cryptococcal meningitis) (Figure 2a, p-value=7.70E-7) as shown in prior studies (51–53), and observed similar outcome ratios across lineages (log-rank test, p-value=0.916) (Figure 2b-c), suggesting no major lineage-specific differences in virulence, though low numbers of VNB and VNII infecting isolates may limit our power to detect significant differences here. Additionally, we noted similar levels of baseline fungal burden and rates of clearance (EFA) between VNIa and VNIb infecting isolates (Figure 2d-e). The data suggest a reduction in baseline fungal burden of 1.29E+06 log_10_ CSF CFU/mL on average for VNII when compared to VNI isolates (Wilcoxon test, p-value=0.024); however, due to the limited number of VNII isolates included, this finding should be confirmed with additional cases.

**Figure 2.**
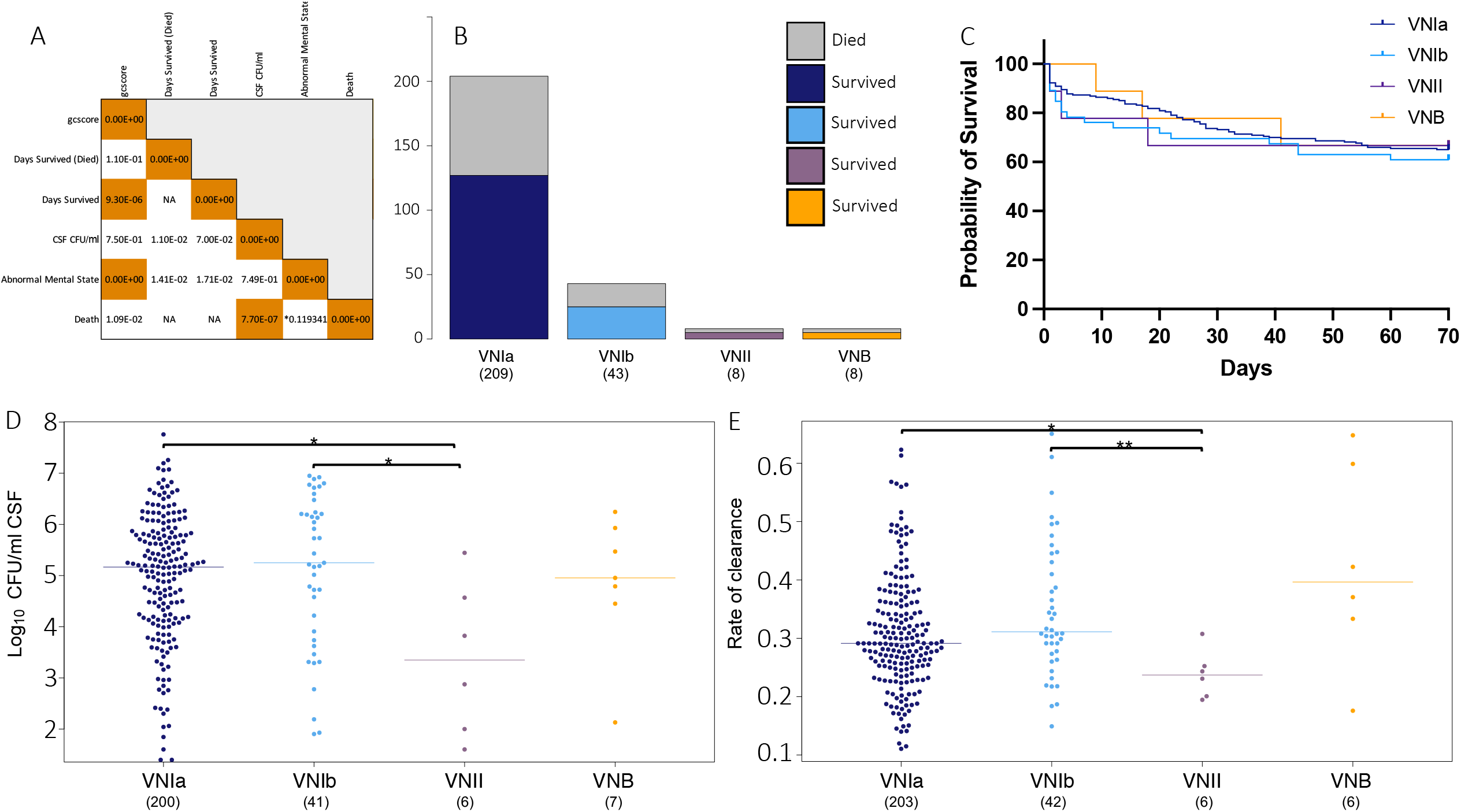
Clinical outcomes by lineage. A) Correlation coefficient between clinical phenotypes, p < 0.0001 in orange. Asterisk indicates phi coefficient. Days survived indicates the total number of days survived for all individuals. Days survived (Died) indicates the number of days survived only for individuals that died during the ACTA clinical trial. B) Deaths (top, grey) and survival (bottom) of patients by infecting isolate lineage. C) Probability of survival, by lineage of infecting isolate. D) CSF Log_10_ CFU/mL (fungal burden) by infecting isolate lineage, asterisk indicates p < 0.05, Wilcoxon test. E) Rate of clearance (EFA) by infecting isolate lineage. Displayed as −1(gradient), asterisk indicates p < 0.05 and 0.01, Wilcoxon test.

To determine whether the variation in baseline fungal burden, which appears well distributed throughout this population (Figure 1), is linked to a specific genetic component, we performed genome-wide association studies (GWAS) to identify variants associated with higher fungal load, when taken as a continuous phenotype. We selected VNI isolates for this analysis as they represent the major genetic group present, and to avoid confounding factors of population structure between lineages. This analysis revealed 53 variants that were significantly (GEMMA score test, p-value < 1.00E-6) associated with CSF fungal burden levels (Figure 3a), 16 of which were predicted to result in a loss-of-function mutation. These variants impacted genes encoding 15 hypothetical proteins and 6 ncRNAs; an additional 4 variants fell into noncoding centromeric regions (Supplemental Table 4). Of the annotated genes impacted, 5 have been previously identified to modulate virulence phenotypes, these include the SAGA histone acetylation complex *SGF29* (CNAG_06392), the protein S-acyl transferase *PFA4* (CNAG_03981), the calcineurin catalytic subunit *CNA1* (CNAG_04796), and the mitochondrial co-chaperone *MRJ1* (CNAG_00938) that are required for virulence in the murine model, as well as the iron permease *FTR1* (CNAG_06242) that is required for capsule regulation (54–57). Additionally, 2 genes with variants are known to impact titan cell formation, these include the multidrug transporter CNAG_04546 and the adenylate cyclase *CAC1* (CNAG_03202) (58, 59). A high proportion (28%) of variants with high GWAS scores (GEMMA score test, p < 1.00E-6) appeared in genes annotated as hypothetical proteins. We additionally performed GWAS to associate variants with mortality due to the correlation between mortality and high fungal burden. However, we did not observe any variants significantly associated with mortality, perhaps due to the multiple host factors that are known to impact this outcome, including the immune status of the host, raised intracranial pressure, duration of infection, and toxicities/adverse effects of antifungal drugs. For phenotype characterization, we decided to focus on genes impacted by variants associated with higher fungal burden, that were predicted to result in a loss-of-function or amino acid change within coding regions.

**Figure 3.**
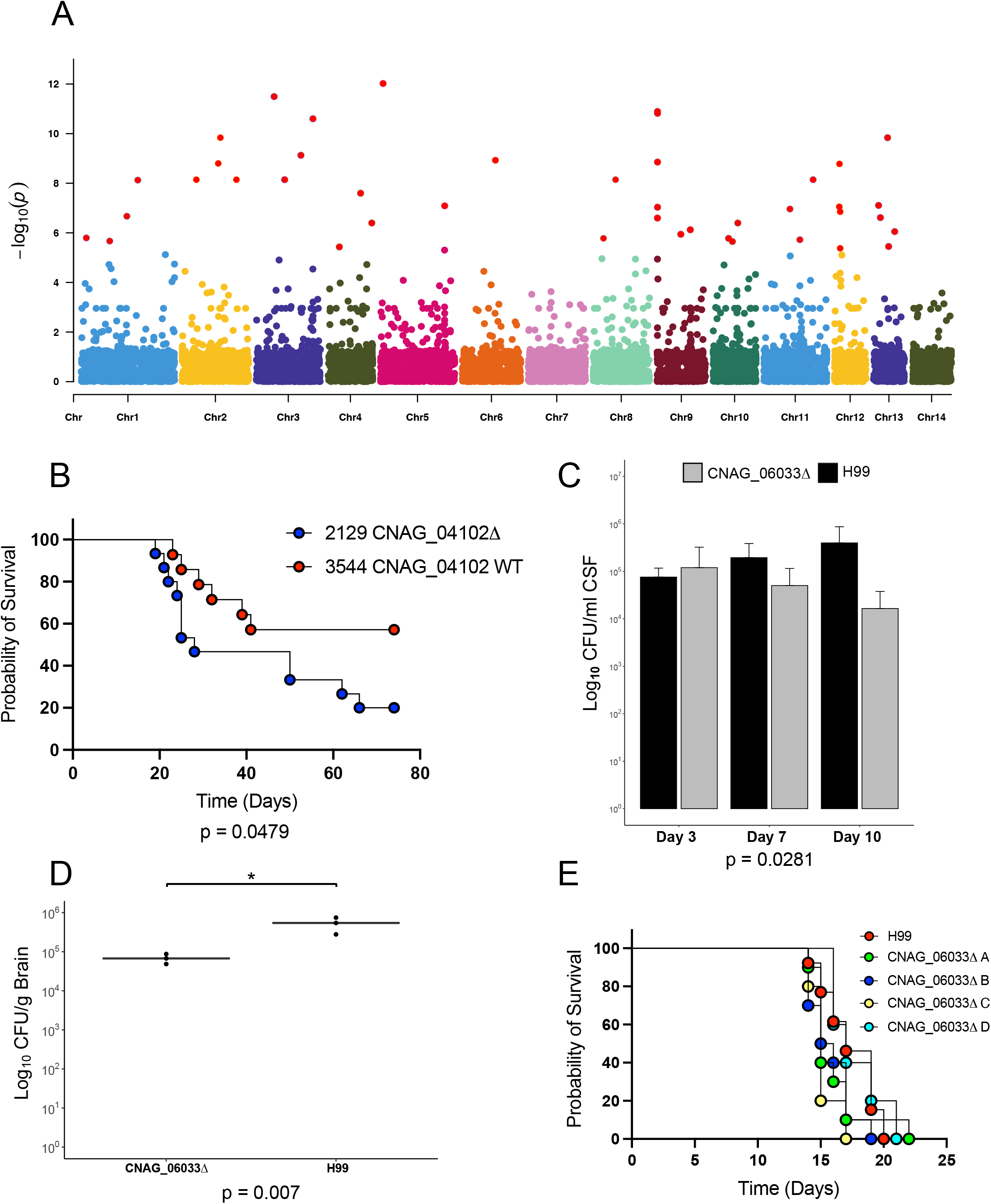
Impact on virulence of genes containing variants significantly associated with fungal burden. A) Manhattan plot displaying variants associated with high fungal burden. Variants with an association score < 0.000005 (score test) are labeled in red. B) Survival of mice infected with closely related ACTA isolates 2129 and 3544. 15 CD-1 mice were infected with approximately 5×10^4^ CFU by oropharyngeal aspiration. C) Rabbit CSF CFU’s for the parental strain (H99) and two independent CNAG_06033 mutant strains extracted on days 3, 7, and 10 post-infection. Three rabbits were infected per strain, n=3 per strain at days 3 and 7, n=2 per strain at day 10. D) Rabbit Log_10_ CFU/g for parental (H99) and one CNAG_06033 mutant strain, extracted from brain tissue (right lobe) at 10 days post-infection. E) Survival of mice infected with parental strain (H99) and four independent CNAG_06033 mutant strains. Five CD-1 mice were infected with approximately 5×10^4^ CFU by oropharyngeal aspiration.

### Hypothetical proteins impact virulence in a strain and background dependent manner

To determine if the genes identified through our GWAS analysis of fungal burden impact virulence in animal models, we tested a total of 10 gene deletion strains across murine and rabbit models. Previous work has shown that infection outcomes from human infections can be recapitulated in murine models (8). Furthermore, rabbit models have proven useful in evaluating CNS infections, as fungal burden within the CNS can be determined through longitudinal collection of CSF (60). The most striking result from our GWAS analysis was a set of 5 different variants in the same hypothetical protein, CNAG_04102, with the highest scoring frameshift variant within this gene being the third most significant overall (GEMMA score test, p=1.30E-11). There were 12 isolates with natural loss-of-function variants in CNAG_04102. To determine whether these variants impact virulence, we chose two closely related clinical isolates (ACTA2129 and ACTA3544), with and without a frameshift in CNAG_04102, and tested them in a murine model. The isolate with a frameshift in CNAG_04102 showed significantly increased rates of mortality (p=0.0479) (Figure 3b), when compared to the control isolate. However, when two independent deletions of this gene were generated in a H99 background and tested in a murine model, we observed no significant impact on virulence (Supplemental Figure 6), highlighting the complex impact of these variants in a background specific manner. CNAG_04102 contains a Kyakuja-Dileera-Zisupton (KDZ) superfamily motif (pfam18758), which has been found within species from basidiomycota, mucoromycotina, *Rhizophagus* and *Allomyces* (61), with CNAG_04102 homologs containing this motif found in *Cryptococcus gattii* and *Cryptococcus floricola*. The KDZ motif is also commonly located within TET/JBP genes which are involved in genomic organization and epigenetic regulation (62), suggesting a role for gene expression regulation. A second hypothetical protein, CNAG_05608, displayed missense variants associated with fungal burden in 22 isolates. An available CNAG_05608 deletion strain in the CMO26 KN99 background (38) showed reduced virulence within a murine model, when compared to wild type CMO26 (log-rank test, p=0.0154) (Supplemental Figure 6). However, when two independent deletions of CNAG_05608 were tested in a H99 background, we saw no difference in virulence, again highlighting the impact of strain background in virulence phenotypes (Supplemental Figure 6). While CNAG_05608 is annotated as a hypothetical protein, this gene is predicted to contain a single transmembrane domain and has homologs in *Cryptococcus gattii*, *Cryptococcus amylolentus*, *Kwoniella* species, and *Wallemia* species. Furthermore, this gene is upregulated during growth in both murine and monkey lungs (63), and slightly downregulated when grown in the presence of glucose (64), suggesting a role during infection.

### Sugar transport and metabolism impacts persistence in a rabbit CNS infection model

In testing for the association of loss-of-function variants with fungal burden, the most highly significant variant was a frameshift in the phosphofructokinase gene, CNAG_06033 (pfkB) (GEMMA score test, p=1.69E-09). The deletion mutants resulting from four independent deletions of CNAG_06033 in a H99 background displayed a trend towards increased virulence in a murine model when compared to H99 (17 day median survival for H99, 15.5 day median survival for CNAG_06033 deletions), though not statistically significant (log-rank test, p=0.0637) (Figure 3). Two mice with CFU counts for H99 were excluded at day 42 as they were identified as outliers via ROUT analysis. Paradoxically, two independent deletion mutants of this gene displayed significantly decreased CSF burden within the rabbit model (3 New Zealand white male rabbits) when compared to the parental strain, with CSF levels (log_10_ CFU/mL counts) dependent on both the infecting strain and the number of days post-infection. The CSF loads were comparable across three rabbits for H99 and the two CNAG_06033 deletion strains at 3 days post-infection but decreased significantly for the CNAG_06033 deletion strains at days 7 and 10, in contrast to H99 which showed increased CSF load over time (repeated measures analysis, p=0.028) (Figure 3c). This highlights the need for efficient glycolysis within the mammalian CSF, as the disruption of both early and late glycolysis regulatory genes, phosphofructokinase (CNAG_06033) and pyruvate kinase (45), reduces *C. neoformans* growth within the rabbit CSF. Congruently, the fungal burden within the brain of rabbits infected with a CNAG_06033 deletion strain also appeared significantly reduced when compared to H99 (T-test, p=0.007) (Figure 3d). Loss-of-function variants within CNAG_06033 have been shown to emerge over the course of in vivo passage in CSF during human infection and relapse (15), suggesting a role for the loss of CNAG_06033 in adaptation to a specific host. However, in an acute rabbit infection model, the loss of CNAG_06033 results in lower survival for *C. neoformans*. This result is consistent with prior work that tested deletions of other genes directly involved in glycolysis; loss of pyruvate kinase (*pyk1Δ*) resulted in decreased persistence in the rabbit CSF, but unperturbed dissemination in a murine model (45). To better understand the requirement of glycolysis and sugar transport in the efficient survival of *C. neoformans* within the CSF, we identified significant variants in additional genes involved in these pathways. A predicted xylose transporter, CNAG_05324, contained a frameshift variant present in 33 isolates (GEMMA score test, p=4.00E-07). In preliminary experiments deletion of CNAG_05324 in a H99 background led to an increase in CSF load in one rabbit when compared to its H99 control. However, additional experiments are required to confirm these results (Supplemental Figure 4a). Given the predicted role of this gene as a xylose transporter, and the presence of xylose in cryptococcal capsule, we undertook preliminary phenotypic capsule screening of the CNAG_05324 deletion strain. Capsule analysis of this deletion mutant via cultivation in capsule-inducing media and India ink staining revealed a significant increase in capsule thickness (Welch’s T-test, p=3.9E-14) (Supplemental Figure 4b), suggesting a role for CNAG_05324 in capsule size modulation. Previous work has shown that modulation of xylose transport and xylosylation can drastically alter virulence, capsule size, and immune evasion (65, 66), highlighting this capsular mechanism as an area for further exploration.

### Aneuploidy is common and slows growth

To determine how natural variation might affect growth and other clinically relevant phenotypes, we performed in vitro phenotyping of isolates. Isolates displayed a range of growth levels on rich media (YPD) at 30°C, 37°C, and 39°C (Figure 4a-c), with colony size across conditions and replicates showing strong correlation (Figure 4d,e, replicate per condition R^2^ > 0.8). To determine whether this variation is linked to a specific genetic component, we performed GWAS to identify variants associated with increased and decreased colony size. Significantly associated with the rapid growth of large colonies on YPD were loss-of-function variants in CNAG_06637 (UBP8 Ubiquitinspecific protease, component of the SAGA complex), CNAG_03818 (sensory transduction histidine kinase), and CNAG_10082 (tRNA Threonine) (GEMMA score test, p < 1.00E-6). A single loss-of-function variant was found significantly associated with decreased colony size, a frameshift in the dolichyldiphosphatase encoding gene, CNAG_03014 (GEMMA score test, p=9.90E-12). Naturally occurring loss-of-function variants such as these may play a role in the fitness variation we see between clinical isolates.

**Figure 4.**
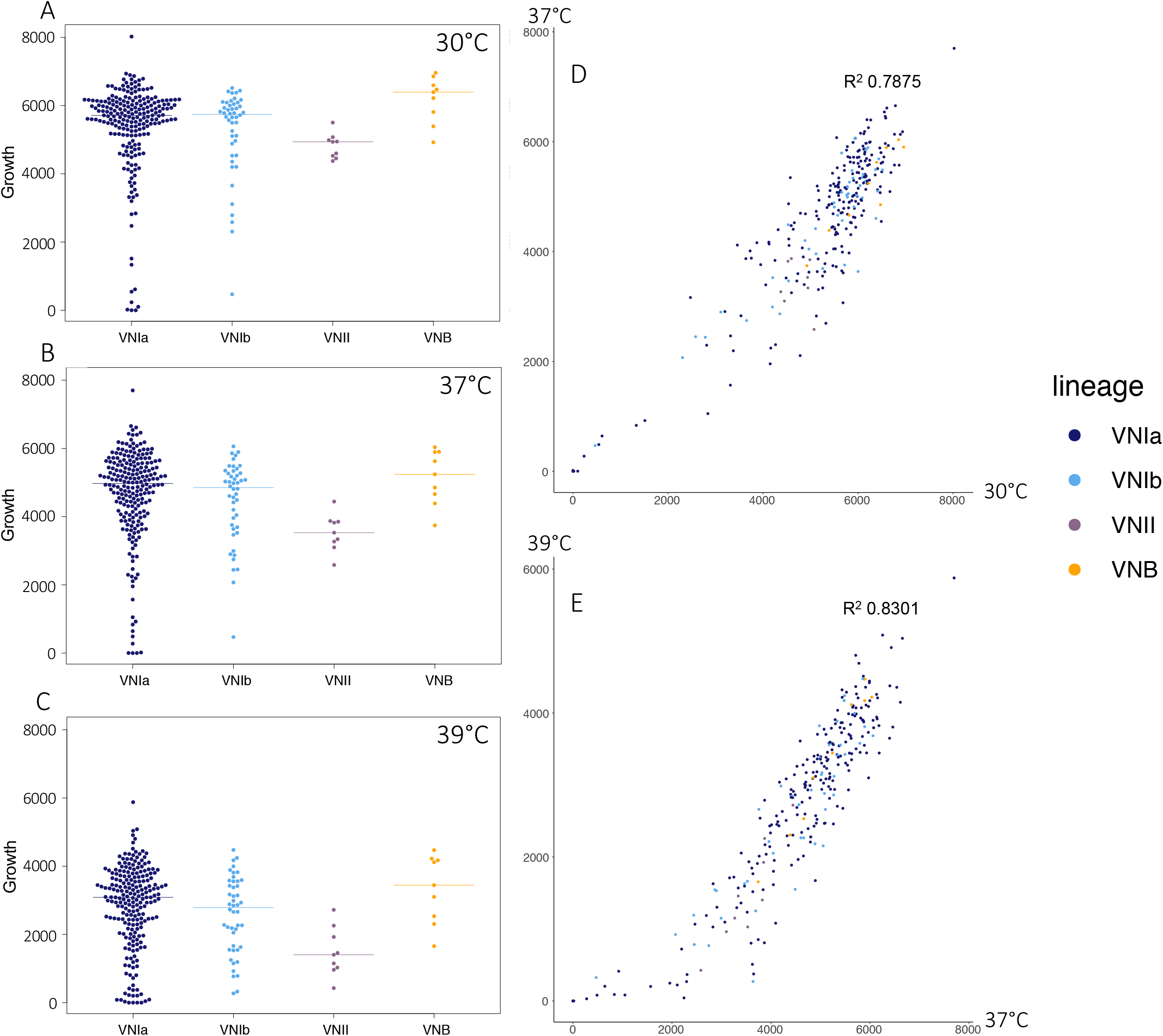
Isolate growth and phenotype correlation. Colony size (area, px) of isolates grown on YPD at A) 30°C, B) 37°C, C) 39°C. Correlation of isolate growth (area, px) on YPD, with axes corresponding to colony size when grown at D) 30°C and 37°C, E) 37°C and 39°C. Colors correspond to isolate lineage: VNIa (dark blue), VNIb (light blue), VNII (purple), VNB (orange).

In addition to SNP and indel mutations, we evaluated the level of chromosome copy number variation across these clinical isolates. We observed both fully and partially duplicated chromosomes, with the most commonly duplicated chromosomes being 12, 9, 14, and 1; overall, duplication of an entire chromosome occurs in 8.5% of clinical isolates. Partial duplications, where at least 25% of the chromosome shows continuous duplication, occur most frequently in chromosomes 2 and 6 (15 instances, Figure 5a). Aneuploid isolates appear well distributed throughout this clinical dataset (Supplemental Figure 5a), suggesting frequent and independent origins for these events occurring in vivo.

**Figure 5.**
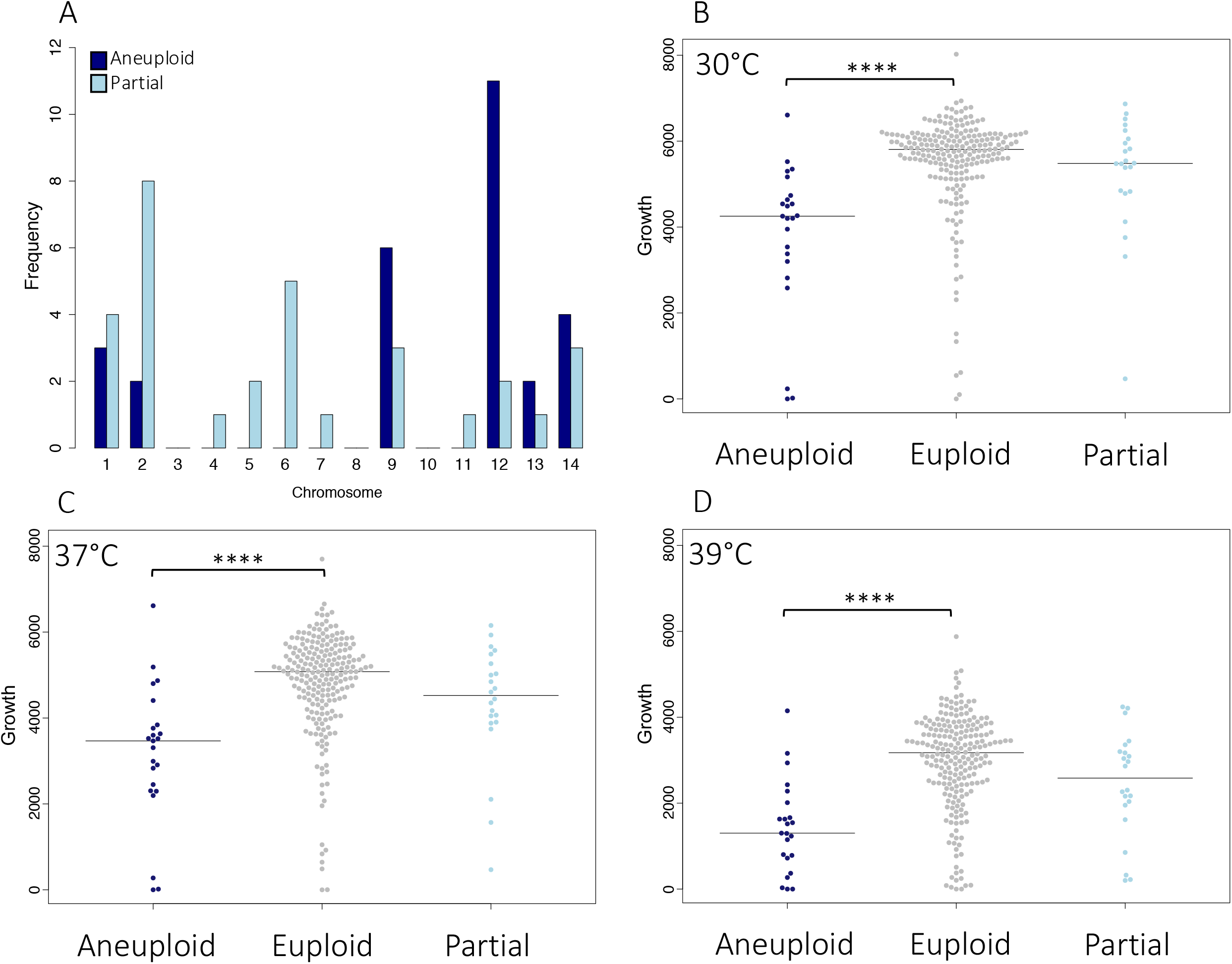
Impact of aneuploidy on growth phenotypes. A) Whole (aneuploid) and partial chromosomal duplication frequency throughout the population, by chromosome. Colony size (growth) on YPD, by ploidy state at B) 30°C, C) 37°C, D) 39°C.

To evaluate the impact of these large aneuploidies, we next compared the growth of aneuploid and euploid isolates. As aneuploidy, specifically of chromosome 1, has been linked to azole resistance, we first assessed the ability of these aneuploid isolates to grow in the presence of fluconazole at 64 μg/mL. After 3 days, we observed that 4% (2 isolates, of which 1 showed a chromosome 1 aneuploidy) of the aneuploid isolates were capable of growth under this condition, while 9% (22 isolates) of non-aneuploid isolates were capable of growth on fluconazole. As aneuploidy across a range of chromosomes did not appear to offer a specific advantage to growth on fluconazole, we went on to assess the growth phenotypes of these isolates at a range of temperatures. On rich media (YPD), at 30°C, 37°C, and 39°C, isolates harboring a fully duplicated chromosome showed significantly poorer growth than euploid isolates (Wilcoxon test, p < 5.00E-07), or isolates featuring only a partial chromosomal duplication (Wilcoxon test, p < 0.01) (Figure 5b-d). To determine whether this fitness reduction occurs in both clinical and environmental populations, we carried out a metanalysis of the data for these isolates, and data previously generated using the same assay for a diverse set of clinical and environmental isolates (Supplemental Figure 5b) (11). Isolates with aneuploidies present in both datasets displayed a significant reduction in growth on YPD at 37°C (p=2.00E-07), demonstrating that this reduction in fitness holds true for both clinical and environmental isolates, across lineages VNI and VNB. We did not observe any significant association between aneuploidy and any single clinical factor; however, we did note that aneuploid isolates showed a reduced fungal burden in patients (825,928 average CFU/mL), when compared to their euploid counterparts (1,318,576 average CFU/mL), though this difference was not statistically significant.

Aneuploidy of specific chromosomes may be advantageous under certain stressors such as antifungal treatment; however, with optimal nutrients at a range of temperatures we found that aneuploidy significantly reduces fitness. Modulation of chromosome 1 ploidy has been linked to apoptosis-inducing factor 1 (*AIF1*) (13); however, variants within *AIF1* were not present in this population, suggesting an alternate mechanism may be responsible for the modulation of chromosome copy number here.

## Discussion

Phylogenetic analysis of 284 *C. neoformans* samples from patients with cryptococcosis in Malawi revealed a mixed population dominated by the VNI genetic group, the most commonly observed global lineage of *C. neoformans*. This population consists of two of the three previously identified VNI lineages (VNIa, and VNIb) (67, 68). VNI isolates are found around the globe and appear relatively clonal when compared to the highly diverse VNB lineage that is often isolated from rural niches, such as mopane trees, in Africa and South America (11, 69–71).

Sugar transporters have previously been identified as under selection in both VNI and VNB isolates from Botswana (11), and we find they also appear under selection in both VNI and VNB isolates in this population from Malawi. Selection pressures in *Cryptococcus* likely occur in the environment and provide a coincidental advantage during human infection; for example, transporter gene evolution has been proposed to be driven in part by adaptation to different trees (72). We find inositol, xylose, glucose, lactose, and glucoside transporters under selection. The expansion of inositol transporters in *C. neoformans* may offer an advantage in both woodland areas and the CNS, as these environments have abundant inositol (47). Xylose transport is important for *C. neoformans* capsule production and the variable xylosylation of an isolate may enable immune evasion (66). The signaling molecule and preferred carbon source glucose, the precursor of glucoside, is known to regulate hexose transporters that are required for virulence (48) and is a key glycolytic metabolite, a pathway required for growth in the CNS (45).

We identified multiple variants significantly associated with clinical and growth phenotypes by taking a GWAS approach; however, clinical phenotypes such as mortality and mental-status did not show a strong association with any variants identified, perhaps due to the complex nature of these clinical characteristics. Additionally, we used a culture-based method to select for isolates from patients, and culture-negative individuals were thus excluded. As a result, we were unable to detect variants associated with very low levels of fungal burden in the CNS. While high fungal burden and mortality are correlated within this dataset, additional host factors such as immune responses are also known to modulate patient outcome (73, 74). In this cohort, while CSF burden has a high correlation with infection outcome, there are multiple cases in which the individual either survived or died despite very high or low initial fungal burden respectively. These confounding factors may explain why we see a strong association of variants with fungal burden, but not with more complex phenotypes such as mortality. While extensively applied to human data, genome-wide association studies have also been applied to study fungal human pathogens (9–11) and plant-pathogens (75, 76). A major challenge is adapting these association approaches for the population structure of each species; for *C. neoformans*, while there is recombination within the population, LD50 values for VNI, VNBI, and VNBII populations are < 50kb (11). Expanding the sample size for such associations or focusing narrowly on particular genetic groups can help increase the power to detect variants; however, the choice of GWAS approach also needs to be optimized for the population under study through consideration of population structure and size.

Through analysis of the clinical metadata, we found CSF fungal burden, a measure of the quantity of live yeasts at the site of infection, to be the most strongly associating clinical parameter in carrying out GWAS. We found that high fungal burden within the CSF of an individual strongly correlated with patient mortality, in accordance with prior work showing that high fungal load is a predictor of mortality (51–53). Multiple variants were associated with high fungal burden and when a pair of clinical isolates were tested in a murine model, the isolate containing a frameshift in CNAG_04102 showed increased virulence, when compared to its wild-type counterpart. Conversely, when we tested two independent deletions of CNAG_04102 in a H99 background, we did not see this phenotype recapitulated, suggesting other background specific factors may be at play. Similarly, when we tested an available deletion of another gene impacted by these variants, CNAG_05608, in the KN99 background, we saw reduced virulence in a mouse infection model; however, this was not phenocopied in the H99 background. Furthermore, the variants significantly associated with high fungal burden in CNAG_05608 were exclusively missense variants, which may have altered the protein’s functionality in a way that did not result in such a drastic loss of function as a deletion strain. Of note, when records were analyzed for the individuals infected with isolates containing missense variants in CNAG_05608, we found that a similar proportion of these patients died when compared to overall mortality within this cohort (odds ratio=1.591, p-value=0.431).

We found that isolates lacking a functional phosphofructokinase B (CNAG_06033) exhibited slightly accelerated mortality in mice but a reduced CSF load within a rabbit model. We hypothesize that these conflicting findings reveal how *Cryptococcus* adjust its energy use during infection in different hosts. Phosphofructokinase plays a key regulatory role in the glycolytic pathway, which appears to be required for optimal virulence in the rabbit model. However, loss-of-function variants in CNAG_06033 were associated with higher fungal burden in patients and reduced survival in mice. This later phenotype is consistent with the hypervirulence phenotype previously observed when comparing serial isolates from the same patient; a persistent isolate harboring a frameshift in CNAG_06033 showed hypervirulence in a galleria model, when compared to the initial infecting isolate (15). Temporal as well as environmental differences between the human and rabbit CNS may alter the expression of, and reliance on, specific metabolic pathways in these two environments. For instance, CNAG_06033 is differentially expressed when comparing *Cryptococcus* samples taken from human CSF, where it is downregulated, with those from rabbit CSF, where it is upregulated. It is also differentially expressed across the course of infection in the rabbit infection model, when comparing days 1 and 4 post-infection in the rabbit CSF. When comparing the cryptococcal transcription of genes in metabolic pathways between rabbit and human CSF samples, those involved in glycogen metabolism appear significantly upregulated in human CSF when compared to rabbit CSF, this observation suggests an increased reliance on the metabolism of stored carbon sources for the yeast in the human host (23), hinting that efficient glycolysis may be less necessary for cryptococcal growth in this setting. Within this ACTA cohort, patient isolates containing naturally occurring loss-of-function variants in CNAG_06033 showed similar mortality when compared to overall mortality (odds ratio=0.774, p-value=0.770). These isolates showed no growth defects when grown on YPD at 37°C, and our phosphofructokinase mutant showed a CSF load reduction similar to that observed for the pyruvate kinase (*PYK1*) deletion strain within a rabbit model, likely due to the similar regulatory effects of both enzymes in glycolysis in an acute rabbit model of infection (45). Additionally, loss-of-function variants have previously been identified in CNAG_06033 (PfkB) after in vivo human passage (15). Under stress conditions, metabolically heterogeneous yeast populations may emerge (77, 78), this metabolic diversity might explain the emergence of isolates less reliant upon glycolysis for chronic growth within the host, as shutting down a major regulatory gene of glycolysis may allow for more efficient gluconeogenesis, use of fatty acids, and beta oxidation for energy (79). In contrast, for growth within the acute rabbit model, *Cryptococcus* relies upon an intact glycolytic pathway for efficient stress response and growth in the CNS. These results demonstrate the importance of both body site and timing for central metabolic pathways during infection.

This unbiased GWAS approach has allowed us to identify hypothetical proteins implicated in virulence, in the absence of additional functional or pathway information. Other systematic studies utilizing RNA-Seq have also identified genes encoding hypothetical proteins that are strong candidates for further study, due to their high expression in CSF (23, 46). While the large proportion of *C. neoformans* genes annotated as hypothetical proteins are more challenging to study, it is critical that we more widely characterize the roles of all genes involved in the pathogenesis of *C. neoformans*.

We found evidence that ploidy directly impacts the fitness of both clinical and environmental *C. neoformans* isolates. Aneuploidy has been linked to broad-spectrum stress resistance in *Candida* species (80), and in *C. neoformans*, disomy of chromosome 1 is known to arise in isolates treated with azoles both in vitro and in vivo and confers resistance to azoles such as fluconazole, through the increase in copy number of *AFR1* and *ERG11* (12, 14). Suggested mechanisms for modulation of chromosome 1 ploidy include regulation via the apoptosis-inducing factor Aif1 (13); however, we did not find evidence for Aif1 disruption in these isolates. Specific impacts of disomy have also been noted for chromosome 13, disomy of which results in reduced melanization (20). In *S. cerevisiae*, disomy of select chromosomes also results in reduced proliferation rates (81). Whilst the reduction in fitness we observed does not seem specific to any particular chromosome, the questions of how and why ploidy appears subject to change in stressful conditions and whether the most frequently observed aneuploidies confer a specific advantage are intriguing and will require further study.

By combining genetic, in vitro, clinical data, and animal model validation, we gain insights into the impact of naturally-occurring genetic variation and its implication for infection outcomes. As whole genome sequencing on an ever-larger scale becomes more accessible, and association techniques for fungal populations grow in sophistication, so too will our power to detect functionally relevant genetic variation across cryptococcal populations. Combined with data from large pan-African clinical trials, and approaches that leverage both fungal and human variant identification, we can further dissect the interactions between pathogen and host genetics. Together, this will enable a better understanding of how these variations impact the ability of *Cryptococcus* to adapt to, and thrive in, the wide range of environments it finds itself in.

## Data Availability

Isolate sequence data can be accessed in NCBI via accession PRJNA764746.

## Ethics

The ACTA trial from which the isolates described here were collected had ethical approval from the London School of Hygiene and Tropical Medicine Research Ethics Committee and by all the site national research ethics committees and regulatory bodies. De-identified clinical metadata (fungal burden, fungal clearance (EFA), patient outcome) was provided by investigators for analysis performed here.

## Acknowledgments

We thank the Broad Institute Genomics Platform for generating the sequence data for this study. We thank all of the patients and their families, as well as the staff at all the sites not directly involved in the ACTA trial; Andrew Nunn, Halima Dawood, Andrew Kitua, and William Powderly for serving on the data and safety monitoring committee; Graeme Meintjes, Calice Talom, Newton Kumwenda, and Maryline Bonnet for serving on the trial steering committee; and the ANRS Staff in Paris (Brigitte Bazin, Claire Rekacewicz, and Paula Garcia) for constant support.

## Figure Legends

Supplemental Figure 1. Linkage disequilibrium decay over 250kb for lineages VNIa, VNIb, and VNI (VNIa + VNIb). VNI shows 50% LD decay in 30kb.

Supplemental Figure 2. Regions under selection per lineage, determined by CLR analysis. Highlighted regions are in the top 5% (orange) or top 10% (green) of all regions. Centromeric regions are highlighted in light grey. Region enrichment via hypergeometric testing for GO categories by lineage, with contributing genes and adjusted p-values given.

Supplemental Figure 3. Growth on fluconazole by lineage. Colony size (area, px) of isolates grown on fluconazole 64 μg/mL for 3 days.

Supplemental Figure 4. Capsule and virulence regulation of CNAG_05324. A) Rabbit CSF load, Log_10_(CFU/mL), on days 3, 7, and 10, per rabbit (individual lines) infected with either H99 (blue) or the CNAG_05324 deletion strain (pink). B) Cell size and capsule diameter at 48 hours growth in capsule inducing media for H99 and the CNAG_05324 deletion strain.

Supplemental Figure 5. Aneuploidy impact on growth for ACTA and Desjardins et al. isolates. A) Whole (aneuploid) and partial chromosomal duplications throughout the population. B) Colony size (growth) by ploidy state on YPD at 37°C for both ACTA and Desjardins et al isolates.

Supplemental Figure 6. Survival of mice infected with parental strain (H99/CMO26) or deletion strain. Five CD-1 mice were infected with approximately 5×10^4^ CFU by oropharyngeal aspiration per strain. A) Survival of mice infected with parental strain (H99) and a CNAG_00544 mutant strain. B) Survival of mice infected with parental strain (CMO26) and a CNAG_02777 mutant strain. C) Survival of mice infected with parental strain (CMO26) and a CNAG_05119 mutant strain. D) Survival of mice infected with parental strain (H99) and two independent CNAG_04102 mutant strains. E) Survival of mice infected with parental strain (CMO26) and a CNAG_04548 mutant strain. F) Survival of mice infected with parental strain (CMO26) and a CNAG_03754 mutant strain. G) Survival of mice infected with parental strain (CMO26) and a CNAG_03161 mutant strain. H) Survival of mice infected with parental strain (H99/CMO26) and three independent CNAG_05608 mutant strains in the CMO26 background (C) and the H99 background (A, B).

Supplemental Table 1. Wild type and mutant strains used in this study.

Supplemental Table 2. Genes under selection for isolates of lineages VNI, VNB and VNII.

Supplemental Table 3. Genes and gene annotations for duplicated regions identified via copy number variation analysis.

Supplemental Table 4. Genes impacted by variants significantly associated with high cerebrospinal fluid fungal burden.

## References

1. Rajasingham R, Smith RM, Park BJ, Jarvis JN, Govender NP, Chiller TM, Denning DW, Loyse A, Boulware DR. 2017. Global burden of disease of HIV-associated cryptococcal meningitis: an updated analysis. Lancet Infect Dis 17:873–881.

2. Maziarz EK, Perfect JR. 2016. Cryptococcosis. Infect Dis Clin North Am 30:179–206.

3. Chayakulkeeree M, Perfect JR. 2006. Cryptococcosis. Infect Dis Clin North Am 20:507–544, v–vi.

4. Litvintseva AP, Thakur R, Vilgalys R, Mitchell TG. 2006. Multilocus sequence typing reveals three genetic subpopulations of *Cryptococcus neoformans* var. *grubii* (serotype A), including a unique population in Botswana. Genetics 172:2223–2238.

5. Chen Y, Litvintseva AP, Frazzitta AE, Haverkamp MR, Wang L, Fang C, Muthoga C, Mitchell TG, Perfect JR. 2015. Comparative analyses of clinical and environmental populations of *Cryptococcus neoformans* in Botswana. Mol Ecol 24:3559–3571.

6. Nyazika TK, Hagen F, Machiridza T, Kutepa M, Masanganise F, Hendrickx M, Boekhout T, Magombei-Majinjiwa T, Siziba N, Chin’ombe N, Mateveke K, Meis JF, Robertson VJ. 2016. *Cryptococcus neoformans* population diversity and clinical outcomes of HIV-associated cryptococcal meningitis patients in Zimbabwe. J Med Microbiol 65:1281–1288.

7. Fernandes KE, Brockway A, Haverkamp M, Cuomo CA, van Ogtrop F, Perfect JR, Carter DA. 2018. Phenotypic variability correlates with clinical outcome in *Cryptococcus* isolates obtained from Botswanan HIV/AIDS patients. mBio 9(5):e02016–18.

8. Mukaremera L, McDonald TR, Nielsen JN, Molenaar CJ, Akampurira A, Schutz C, Taseera K, Muzoora C, Meintjes G, Meya DB, Boulware DR, Nielsen K. 2019. The mouse inhalation model of *Cryptococcus neoformans* infection recapitulates strain virulence in humans and shows that closely related strains can possess differential virulence. Infection and Immunity 87(5):e00046–19.

9. Gerstein AC, Jackson KM, McDonald TR, Wang Y, Lueck BD, Bohjanen S, Smith KD, Akampurira A, Meya DB, Xue C, Boulware DR, Nielsen K. 2019. Identification of pathogen genomic differences that impact human immune response and disease during *Cryptococcus neoformans* infection. mBio 10(4):e01440–19.

10. Beale MA, Sabiiti W, Robertson EJ, Fuentes-Cabrejo KM, O’Hanlon SJ, Jarvis JN, Loyse A, Meintjes G, Harrison TS, May RC, Fisher MC, Bicanic T. 2015. Genotypic diversity Is associated with clinical outcome and phenotype in Cryptococcal meningitis across southern Africa. PLOS Neglected Tropical Diseases 9(6):e0003847.

11. Desjardins CA, Giamberardino C, Sykes SM, Yu C-H, Tenor JL, Chen Y, Yang T, Jones AM, Sun S, Haverkamp MR, Heitman J, Litvintseva AP, Perfect JR, Cuomo CA. 2017. Population genomics and the evolution of virulence in the fungal pathogen *Cryptococcus neoformans*. Genome Research 27:1207–1219.

12. Sionov E, Lee H, Chang YC, Kwon-Chung KJ. 2010. *Cryptococcus neoformans* overcomes stress of azole drugs by formation of disomy in specific multiple chromosomes. PLOS Pathogens 6(4):e1000848.

13. Semighini CP, Averette AF, Perfect JR, Heitman J. 2011. Deletion of *Cryptococcus neoformans* AIF ortholog promotes chromosome aneuploidy and fluconazole-resistance in a metacaspase-independent manner. PLOS Pathogens 7(11):e1002364.

14. Stone NRH, Rhodes J, Fisher MC, Mfinanga S, Kivuyo S, Rugemalila J, Segal ES, Needleman L, Molloy SF, Kwon-Chung J, Harrison TS, Hope W, Berman J, Bicanic T. 2019. Dynamic ploidy changes drive fluconazole resistance in human cryptococcal meningitis. J Clin Invest 129:999–1014.

15. Chen Y, Farrer RA, Giamberardino C, Sakthikumar S, Jones A, Yang T, Tenor JL, Wagih O, Van Wyk M, Govender NP, Mitchell TG, Litvintseva AP, Cuomo CA, Perfect JR. 2017. Microevolution of serial clinical isolates of *Cryptococcus neoformans* var. *grubii* and *C. gattii*. mBio 8(2):e00166–17.

16. Rhodes J, Beale MA, Vanhove M, Jarvis JN, Kannambath S, Simpson JA, Ryan A, Meintjes G, Harrison TS, Fisher MC, Bicanic T. 2017. A population genomics approach to assessing the genetic basis of within-host microevolution underlying recurrent cryptococcal meningitis infection. G3: Genes, Genomes, Genetics 7:1165–1176.

17. Fu MS, Liporagi-Lopes LC, Dos SR, Júnior S, Tenor JL, Perfect JR, Cuomo CA, Casadevall A. 2021. Amoeba predation of *Cryptococcus neoformans* results in pleiotropic changes to traits associated with virulence. mBio 12(2):e00567–21.

18. Gerstein AC, Fu MS, Mukaremera L, Li Z, Ormerod KL, Fraser JA, Berman J, Nielsen K. 2015. Polyploid titan cells produce haploid and aneuploid progeny to promote stress adaptation. mBio 6(5):e01340–15.

19. Berman J. 2016. Ploidy plasticity: a rapid and reversible strategy for adaptation to stress. FEMS Yeast Research 16(3):fow020.

20. Hu G, Wang J, Choi J, Jung WH, Liu I, Litvintseva AP, Bicanic T, Aurora R, Mitchell TG, Perfect JR, Kronstad JW. 2011. Variation in chromosome copy number influences the virulence of *Cryptococcus neoformans* and occurs in isolates from AIDS patients. BMC Genomics 12:526.

21. Ormerod KL, Morrow CA, Chow EWL, Lee IR, Arras SDM, Schirra HJ, Cox GM, Fries BC, Fraser JA. 2013. Comparative genomics of serial isolates of *Cryptococcus neoformans* reveals gene associated with carbon utilization and virulence. G3: Genes, Genomes, Genetics 3:675–686.

22. Rhodes J, Desjardins CA, Sykes SM, Beale MA, Vanhove M, Sakthikumar S, Chen Y, Gujja S, Saif S, Chowdhary A, Lawson DJ, Ponzio V, Colombo AL, Meyer W, Engelthaler DM, Hagen F, Illnait-Zaragozi MT, Alanio A, Vreulink JM, Heitman J, Perfect JR, Litvintseva AP, Bicanic T, Harrison TS, Fisher MC, Cuomo CA. 2017. Tracing genetic exchange and biogeography of *Cryptococcus neoformans* var. *grubii* at the global population level. Genetics 207:327–346.

23. Yu C-H, Sephton-Clark P, Tenor JL, Toffaletti D, Giamberardino C, Haverkamp M, Cuomo C, Perfect J. 2021. Gene expression of diverse *Cryptococcus* isolates during infection of the human central nervous system. mBio 12(6):e0231321.

24. Molloy SF, Kanyama C, Heyderman RS, Loyse A, Kouanfack C, Chanda D, Mfinanga S, Temfack E, Lakhi S, Lesikari S, Chan AK, Stone N, Kalata N, Karunaharan N, Gaskell K, Peirse M, Ellis J, Chawinga C, Lontsi S, Ndong J-G, Bright P, Lupiya D, Chen T, Bradley J, Adams J, van der Horst C, van Oosterhout JJ, Sini V, Mapoure YN, Mwaba P, Bicanic T, Lalloo DG, Wang D, Hosseinipour MC, Lortholary O, Jaffar S, Harrison TS, ACTA Trial Study Team. 2018. Antifungal combinations for treatment of cryptococcal meningitis in Africa. N Engl J Med 378:1004–1017.

25. Fisher S, Barry A, Abreu J, Minie B, Nolan J, Delorey TM, Young G, Fennell TJ, Allen A, Ambrogio L, Berlin AM, Blumenstiel B, Cibulskis K, Friedrich D, Johnson R, Juhn F, Reilly B, Shammas R, Stalker J, Sykes SM, Thompson J, Walsh J, Zimmer A, Zwirko Z, Gabriel S, Nicol R, Nusbaum C. 2011. A scalable, fully automated process for construction of sequence-ready human exome targeted capture libraries. Genome Biology 12(1): R1.

26. Loftus BJ, Fung E, Roncaglia P, Rowley D, Amedeo P, Bruno D, Vamathevan J, Miranda M, Anderson IJ, Fraser JA, Allen JE, Bosdet IE, Brent MR, Chiu R, Doering TL, Donlin MJ, D’Souza CA, Fox DS, Grinberg V, Fu J, Fukushima M, Haas BJ, Huang JC, Janbon G, Jones SJM, Koo HL, Krzywinski MI, Kwon-Chung JK, Lengeler KB, Maiti R, Marra MA, Marra RE, Mathewson CA, Mitchell TG, Pertea M, Riggs FR, Salzberg SL, Schein JE, Shvartsbeyn A, Shin H, Shumway M, Specht CA, Suh BB, Tenney A, Utterback TR, Wickes BL, Wortman JR, Wye NH, Kronstad JW, Lodge JK, Heitman J, Davis RW, Fraser CM, Hyman RW. 2005. The genome of the basidiomycetous yeast and human pathogen *Cryptococcus neoformans*. Science 307:1321–1324.

27. Janbon G, Ormerod KL, Paulet D, Byrnes EJ, Yadav V, Chatterjee G, Mullapudi N, Hon CC, Billmyre RB, Brunel F, Bahn YS, Chen W, Chen Y, Chow EWL, Coppée JY, Floyd-Averette A, Gaillardin C, Gerik KJ, Goldberg J, Gonzalez-Hilarion S, Gujja S, Hamlin JL, Hsueh YP, Ianiri G, Jones S, Kodira CD, Kozubowski L, Lam W, Marra M, Mesner LD, Mieczkowski PA, Moyrand F, Nielsen K, Proux C, Rossignol T, Schein JE, Sun S, Wollschlaeger C, Wood IA, Zeng Q, Neuvéglise C, Newlon CS, Perfect JR, Lodge JK, Idnurm A, Stajich JE, Kronstad JW, Sanyal K, Heitman J, Fraser JA, Cuomo CA, Dietrich FS. 2014. Analysis of the Genome and Transcriptome of *Cryptococcus neoformans* var. *grubii* reveals complex RNA expression and microevolution leading to virulence attenuation. PLoS Genetics 10(4):e1004261.

28. Farrer RA, Desjardins CA, Sakthikumar S, Gujja S, Saif S, Zeng Q, Chen Y, Voelz K, Heitman J, May RC, Fisher MC, Cuomo CA. 2015. Genome evolution and innovation across the four major lineages of *Cryptococcus gattii*. mBio 6(5):e00868–15.

29. Farrer RA, Chang M, Davis MJ, van Dorp L, Yang DH, Shea T, Sewell TR, Meyer W, Balloux F, Edwards HM, Chanda D, Kwenda G, Vanhove M, Chang YC, Cuomo CA, Fisher MC, Kwon-Chung KJ. 2019. A new lineage of *Cryptococcus gattii* (VGV) discovered in the central zambezian miombo woodlands. mBio 10(6):e02306–19.

30. Li H. 2013. Aligning sequence reads, clone sequences and assembly contigs with BWA-MEM. arXiv:13033997 [q-bio].

31. Van der Auwera GA, Carneiro MO, Hartl C, Poplin R, Del Angel G, Levy-Moonshine A, Jordan T, Shakir K, Roazen D, Thibault J, Banks E, Garimella KV, Altshuler D, Gabriel S, DePristo MA. 2013. From FastQ data to high confidence variant calls: the Genome Analysis Toolkit best practices pipeline. Curr Protoc Bioinformatics 43:11.10.1–11.10.33.

32. Cingolani P, Platts A, Wang LL, Coon M, Nguyen T, Wang L, Land SJ, Lu X, Ruden DM. 2012. A program for annotating and predicting the effects of single nucleotide polymorphisms, SnpEff: SNPs in the genome of *Drosophila melanogaster* strain w ^1118^□; iso-2; iso-3. Fly 6:80–92.

33. Stamatakis A. 2014. RAxML version 8: a tool for phylogenetic analysis and post-analysis of large phylogenies. Bioinformatics 30:1312–1313.

34. Abyzov A, Urban AE, Snyder M, Gerstein M. 2011. CNVnator: An approach to discover, genotype, and characterize typical and atypical CNVs from family and population genome sequencing. Genome Res 21:974–984.

35. Pfeifer B, Wittelsbürger U, Ramos-Onsins SE, Lercher MJ. 2014. PopGenome: an efficient Swiss army knife for population genomic analyses in R. Mol Biol Evol 31:1929–1936.

36. Zhou X, Stephens M. 2012. Genome-wide efficient mixed-model analysis for association studies. 7. Nature Genetics 44:821–824.

37. Wagih O, Parts L. 2014. gitter: A robust and accurate method for quantification of colony sizes from plate images. G3 (Bethesda) 4:547–552.

38. Chun CD, Madhani HD. 2010. Applying genetics and molecular biology to the study of the human pathogen *Cryptococcus neoformans*. Methods Enzymol 470:797–831.

39. Idnurm A, Reedy JL, Nussbaum JC, Heitman J. 2004. *Cryptococcus neoformans* virulence gene discovery through insertional mutagenesis. Eukaryot Cell 3:420–429.

40. Toffaletti DL, Rude TH, Johnston SA, Durack DT, Perfect JR. 1993. Gene transfer in *Cryptococcus neoformans* by use of biolistic delivery of DNA. J Bacteriol 175:1405–1411.

41. Zaragoza O, Casadevall A. 2004. Experimental modulation of capsule size in *Cryptococcus neoformans*. Biol Proced Online 6:10–15.

42. National Research Council (US) Committee for the Update of the Guide for the Care and Use of Laboratory Animals. 2011. Guide for the Care and Use of Laboratory Animals, 8th ed. National Academies Press (US), Washington (DC).

43. Nielsen R, Williamson S, Kim Y, Hubisz MJ, Clark AG, Bustamante C. 2005. Genomic scans for selective sweeps using SNP data. Genome Res 15:1566–1575.

44. Chow EWL, Morrow CA, Djordjevic JT, Wood IA, Fraser JA. 2012. Microevolution of *Cryptococcus neoformans* driven by massive tandem gene amplification. Molecular Biology and Evolution 29:1987–2000.

45. Price MS, Betancourt-Quiroz M, Price JL, Toffaletti DL, Vora H, Hu G, Kronstad JW, Perfect JR. 2011. *Cryptococcus neoformans* requires a functional glycolytic pathway for disease but not persistence in the host. mBio 2(3):e00103–e111.

46. Chen Y, Toffaletti DL, Tenor JL, Litvintseva AP, Fang C, Mitchell TG, McDonald TR, Nielsen K, Boulware DR, Bicanic T, Perfect JR. 2014. The *Cryptococcus neoformans* transcriptome at the site of human meningitis. mBio 5(1):e01087–13.

47. Xue C, Liu T, Chen L, Li W, Liu I, Kronstad JW, Seyfang A, Heitman J. 2010. Role of an expanded inositol transporter repertoire in *Cryptococcus neoformans* sexual reproduction and virulence. mBio 1(1):e00084–10.

48. Liu T-B, Wang Y, Baker GM, Fahmy H, Jiang L, Xue C. 2013. The glucose sensorlike protein Hxs1 Is a high-affinity glucose transporter and required for virulence in *Cryptococcus neoformans*. PLOS ONE 8:(5):e64239.

49. Li LX, Rautengarten C, Heazlewood JL, Doering TL. 2018. UDP-Glucuronic acid transport is required for virulence of *Cryptococcus neoformans*. mBio 9(1):e02319–17.

50. Li LX, Rautengarten C, Heazlewood JL, Doering TL. 2018. Xylose donor transport is critical for fungal virulence. PLOS Pathogens 14(1):e1006765.

51. Brouwer AE, Rajanuwong A, Chierakul W, Griffin GE, Larsen RA, White NJ, Harrison TS. 2004. Combination antifungal therapies for HIV-associated cryptococcal meningitis: a randomised trial. Lancet 363:1764–1767.

52. Bicanic T, Muzoora C, Brouwer AE, Meintjes G, Longley N, Taseera K, Rebe K, Loyse A, Jarvis J, Bekker L-G, Wood R, Limmathurotsakul D, Chierakul W, Stepniewska K, White NJ, Jaffar S, Harrison TS. 2009. Independent association between rate of clearance of infection and clinical outcome of HIV-associated cryptococcal meningitis: analysis of a combined cohort of 262 patients. Clin Infect Dis 49:702–709.

53. Jarvis JN, Bicanic T, Loyse A, Namarika D, Jackson A, Nussbaum JC, Longley N, Muzoora C, Phulusa J, Taseera K, Kanyembe C, Wilson D, Hosseinipour MC, Brouwer AE, Limmathurotsakul D, White N, van der Horst C, Wood R, Meintjes G, Bradley J, Jaffar S, Harrison T. 2014. Determinants of mortality in a combined cohort of 501 patients with HIV-associated Cryptococcal meningitis: implications for improving outcomes. Clin Infect Dis 58:736–745.

54. Jung WH, Sham A, Lian T, Singh A, Kosman DJ, Kronstad JW. 2008. Iron source preference and regulation of iron uptake in *Cryptococcus neoformans*. PLOS Pathogens 4(2):e45.

55. Nichols CB, Ost KS, Grogan DP, Pianalto K, Hasan S, Alspaugh JA. 2015. Impact of protein palmitoylation on the virulence potential of *Cryptococcus neoformans*. Eukaryotic Cell 14:626–635.

56. Fu C, Donadio N, Cardenas ME, Heitman J. 2018. Dissecting the roles of the calcineurin pathway in unisexual reproduction, stress responses, and virulence in *Cryptococcus deneoformans*. Genetics 208:639–653.

57. Horianopoulos LC, Hu G, Caza M, Schmitt K, Overby P, Johnson JD, Valerius O, Braus GH, Kronstad JW. 2020. The novel J-Domain protein Mrj1 is required for mitochondrial respiration and virulence in *Cryptococcus neoformans*. mBio 11(3):e01127–20.

58. Okagaki LH, Wang Y, Ballou ER, O’Meara TR, Bahn Y-S, Alspaugh JA, Xue C, Nielsen K. 2011. Cryptococcal titan cell formation is regulated by G-Protein signaling in response to multiple stimuli. Eukaryot Cell 10:1306–1316.

59. Trevijano-Contador N, Oliveira HC de, García-Rodas R, Rossi SA, Llorente I, Zaballos Á, Janbon G, Ariño J, Zaragoza Ó. 2018. *Cryptococcus neoformans* can form titan-like cells in vitro in response to multiple signals. PLOS Pathogens 14(5):e1007007.

60. Perfect JR, Lang SD, Durack DT. 1980. Chronic cryptococcal meningitis: a new experimental model in rabbits. Am J Pathol 101:177–194.

61. Muszewska A, Steczkiewicz K, Stepniewska-Dziubinska M, Ginalski K. 2017. Cut-and-paste transposons in fungi with diverse lifestyles. Genome Biology and Evolution 9:3463–3477.

62. Iyer LM, Zhang D, de Souza RF, Pukkila PJ, Rao A, Aravind L. 2014. Lineagespecific expansions of TET/JBP genes and a new class of DNA transposons shape fungal genomic and epigenetic landscapes. Proc Natl Acad Sci USA 111:1676–1683.

63. Li H, Li Y, Sun T, Du W, Li C, Suo C, Meng Y, Liang Q, Lan T, Zhong M, Yang S, Niu C, Li D, Ding C. 2019. Unveil the transcriptional landscape at the *Cryptococcus*-host axis in mice and nonhuman primates. PLOS Neglected Tropical Diseases 13(7):e0007566.

64. Burke JE, Longhurst AD, Merkurjev D, Sales-Lee J, Rao B, Moresco JJ, Yates JR, Li JJ, Madhani HD. 2018. Spliceosome of a Dynamic RNP at Nucleotide Resolution. Cell 173:1014–1030.e17.

65. Moyrand F, Klaproth B, Himmelreich U, Dromer F, Janbon G. 2002. Isolation and characterization of capsule structure mutant strains of *Cryptococcus neoformans*. Mol Microbiol 45:837–849.

66. Li LX, Hole CR, Rangel-Moreno J, Khader SA, Doering TL. 2020. *Cryptococcus neoformans* evades pulmonary immunity by modulating xylose precursor transport. Infect Immun 88(8):e00288–20.

67. Vanhove M, Beale MA, Rhodes J, Chanda D, Lakhi S, Kwenda G, Molloy S, Karunaharan N, Stone N, Harrison TS, Bicanic T, Fisher MC. 2017. Genomic epidemiology of *Cryptococcus* yeasts identifies adaptation to environmental niches underpinning infection across an African HIV/AIDS cohort. Molecular Ecology 26:1991–2005.

68. Ashton PM, Thanh LT, Trieu PH, Van Anh D, Trinh NM, Beardsley J, Kibengo F, Chierakul W, Dance DAB, Rattanavong S, Davong V, Hung LQ, Chau NVV, Tung NLN, Chan AK, Thwaites GE, Lalloo DG, Anscombe C, Nhat LTH, Perfect J, Dougan G, Baker S, Harris S, Day JN. 2019. Three phylogenetic groups have driven the recent population expansion of *Cryptococcus neoformans*. Nature Communications 10(1):2035.

69. Litvintseva AP, Carbone I, Rossouw J, Thakur R, Govender NP, Mitchell TG. 2011. Evidence that the human pathogenic fungus *Cryptococcus neoformans* var. *grubii* may have evolved in Africa. PLoS ONE 6(5):e19688.

70. Khayhan K, Hagen F, Pan W, Simwami S, Fisher MC, Wahyuningsih R, Chakrabarti A, Chowdhary A, Ikeda R, Taj-Aldeen SJ, Khan Z, Ip M, Imran D, Sjam R, Sriburee P, Liao W, Chaicumpar K, Vuddhakul V, Meyer W, Trilles L, Iersel LJJ van, Meis JF, Klaassen CHW, Boekhout T. 2013. Geographically structured populations of *Cryptococcus neoformans* variety *grubii* in Asia correlate with HIV status and show a clonal population structure. PLOS ONE 8:e72222.

71. Andrade-Silva LE, Ferreira-Paim K, Ferreira TB, Vilas-Boas A, Mora DJ, Manzato VM, Fonseca FM, Buosi K, Andrade-Silva J, Prudente B da S, Araujo NE, Sales-Campos H, Silva MV da, Júnior VR, Meyer W, Silva-Vergara ML. 2018. Genotypic analysis of clinical and environmental *Cryptococcus neoformans* isolates from Brazil reveals the presence of VNB isolates and a correlation with biological factors. PLOS ONE 13(3):e0193237.

72. Xue C. 2012. *Cryptococcus* and Beyond—Inositol utilization and its implications for the emergence of fungal virulence. PLOS Pathogens 8:e1002869.

73. Jarvis JN, Meintjes G, Bicanic T, Buffa V, Hogan L, Mo S, Tomlinson G, Kropf P, Noursadeghi M, Harrison TS. 2015. Cerebrospinal fluid cytokine profiles predict risk of early mortality and immune reconstitution inflammatory syndrome in HIV-associated cryptococcal meningitis. PLoS Pathog 11:e1004754.

74. Siddiqui AA, Brouwer AE, Wuthiekanun V, Jaffar S, Shattock R, Irving D, Sheldon J, Chierakul W, Peacock S, Day N, White NJ, Harrison TS. 2005. IFN-gamma at the site of infection determines rate of clearance of infection in cryptococcal meningitis. J Immunol 174:1746–1750.

75. Palma-Guerrero J, Hall CR, Kowbel D, Welch J, Taylor JW, Brem RB, Glass NL. 2013. Genome wide association identifies novel loci involved in fungal communication. PLoS Genetics 9(8):e1003669.

76. Gao Y, Liu Z, Faris JD, Richards J, Brueggeman RS, Li X, Oliver RP, McDonald BA, Friesen TL. 2016. Validation of genome-wide association studies as a tool to identify virulence factors in *Parastagonospora nodorum*. Phytopathology 106:1177–1185.

77. Alanio A, Vernel-Pauillac F, Sturny-Leclère A, Dromer F. 2015. *Cryptococcus neoformans* host adaptation: toward biological evidence of dormancy. mBio 6(2):e02580–14.

78. Hommel B, Sturny-Leclère A, Volant S, Veluppillai N, Duchateau M, Yu C-H, Hourdel V, Varet H, Matondo M, Perfect JR, Casadevall A, Dromer F, Alanio A. 2019. *Cryptococcus neoformans* resists to drastic conditions by switching to viable but non-culturable cell phenotype. PLOS Pathogens 15(7):e1007945.

79. Kretschmer M, Wang J, Kronstad JW. 2012. Peroxisomal and mitochondrial ß-oxidation pathways influence the virulence of the pathogenic fungus *Cryptococcus neoformans*. Eukaryot Cell 11:1042–1054.

80. Yang F, Teoh F, Tan ASM, Cao Y, Pavelka N, Berman J. 2019. Aneuploidy enables cross-adaptation to unrelated drugs. Molecular Biology and Evolution 36:1768–1782.

81. Torres EM, Sokolsky T, Tucker CM, Chan LY, Boselli M, Dunham MJ, Amon A. 2007. Effects of aneuploidy on cellular physiology and cell division in haploid yeast. Science 317:916–924.

