## Supplemental Figure 1 for "Genomic variation across a clinical *Cryptococcus* population linked to disease outcome"

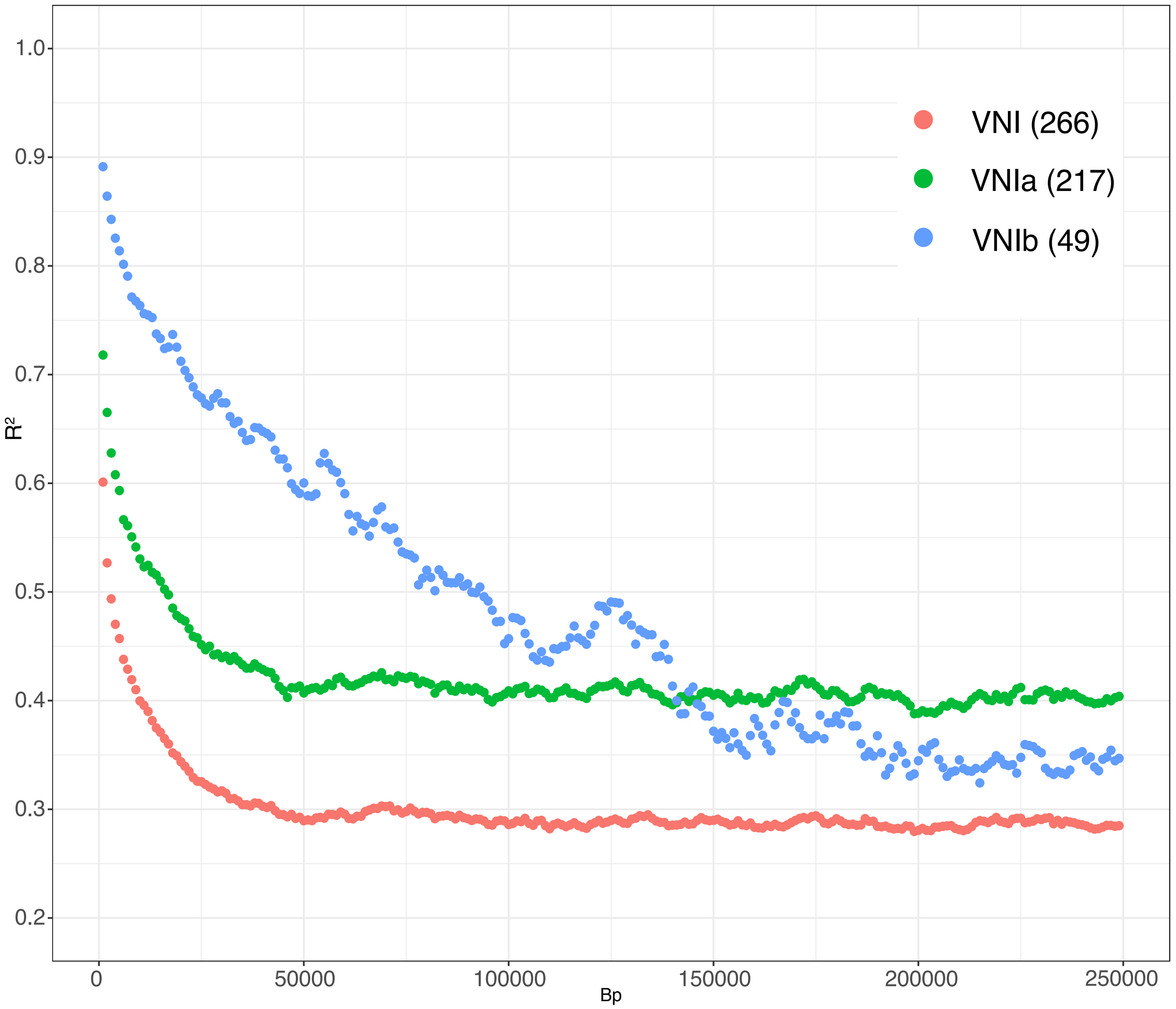


Supplemental Figure 1. Linkage disequilibrium decay over 250kb for lineages VNIa, VNIb, and VNI (VNIa + VNIb). VNI shows 50% LD decay in 30kb.
