## Supplemental Figure 2 for "Genomic variation across a clinical *Cryptococcus* population linked to disease outcome"

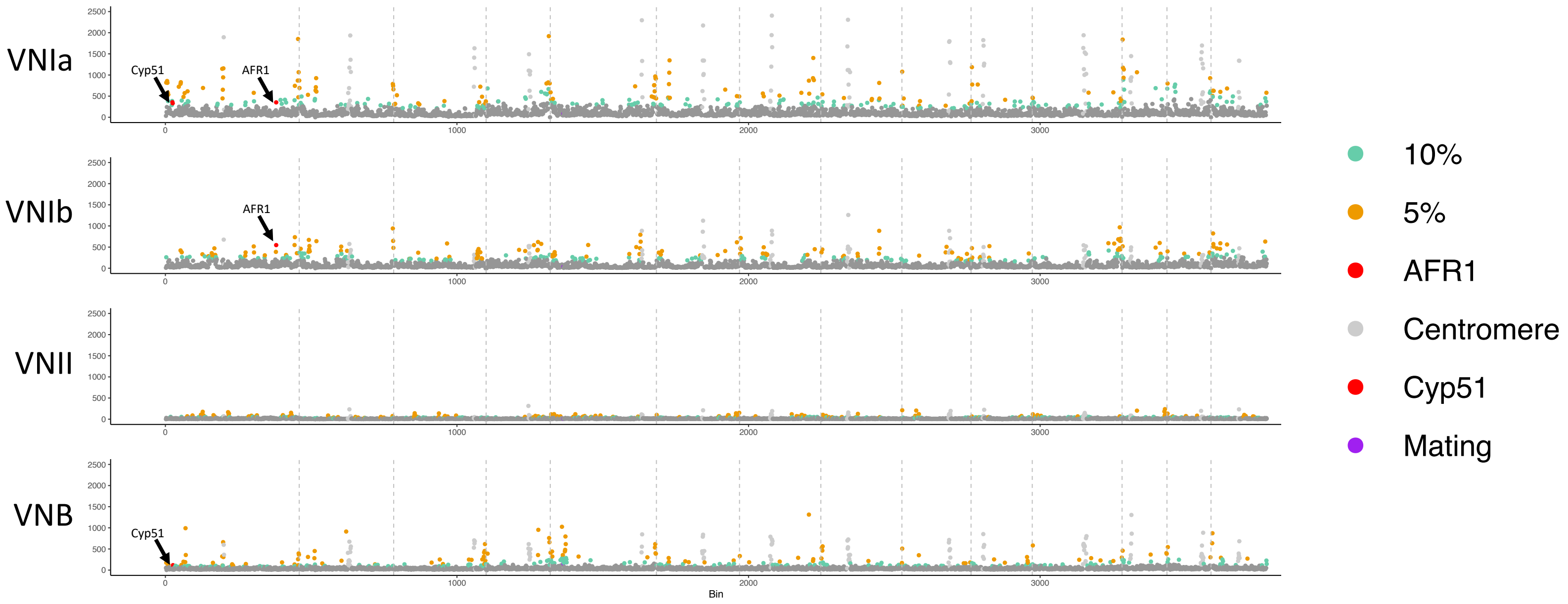


| Lineage | Enriched Category (GO) | Gene Functions | Corrected P Value |
|---|---|---|---|
| VNIa | Carbohydrate Transport | Inositol Transport, Glucose and Glucoside Transport | 1.0765E-3 |
| VNII | Nucleotide Excision Repair | *UVE1, RAD23, RHP42, ERCC2* | 6.7712E-3 |
| VNB | Carbohydrate Transport | Inositol Transport, Glucoside Transport, Galactose Transport | 8.6025E-4 |

Supplemental Figure 2. Regions under selection per lineage, determined by CLR analysis. Highlighted regions are in the top 5% (orange) or top 10% (green) of all regions. Centromeric regions are highlighted in light grey. Region enrichment via hypergeometric testing for GO categories by lineage, with contributing genes and adjusted p-values given.
