## Supplemental Figure 3 for "Genomic variation across a clinical *Cryptococcus* population linked to disease outcome"

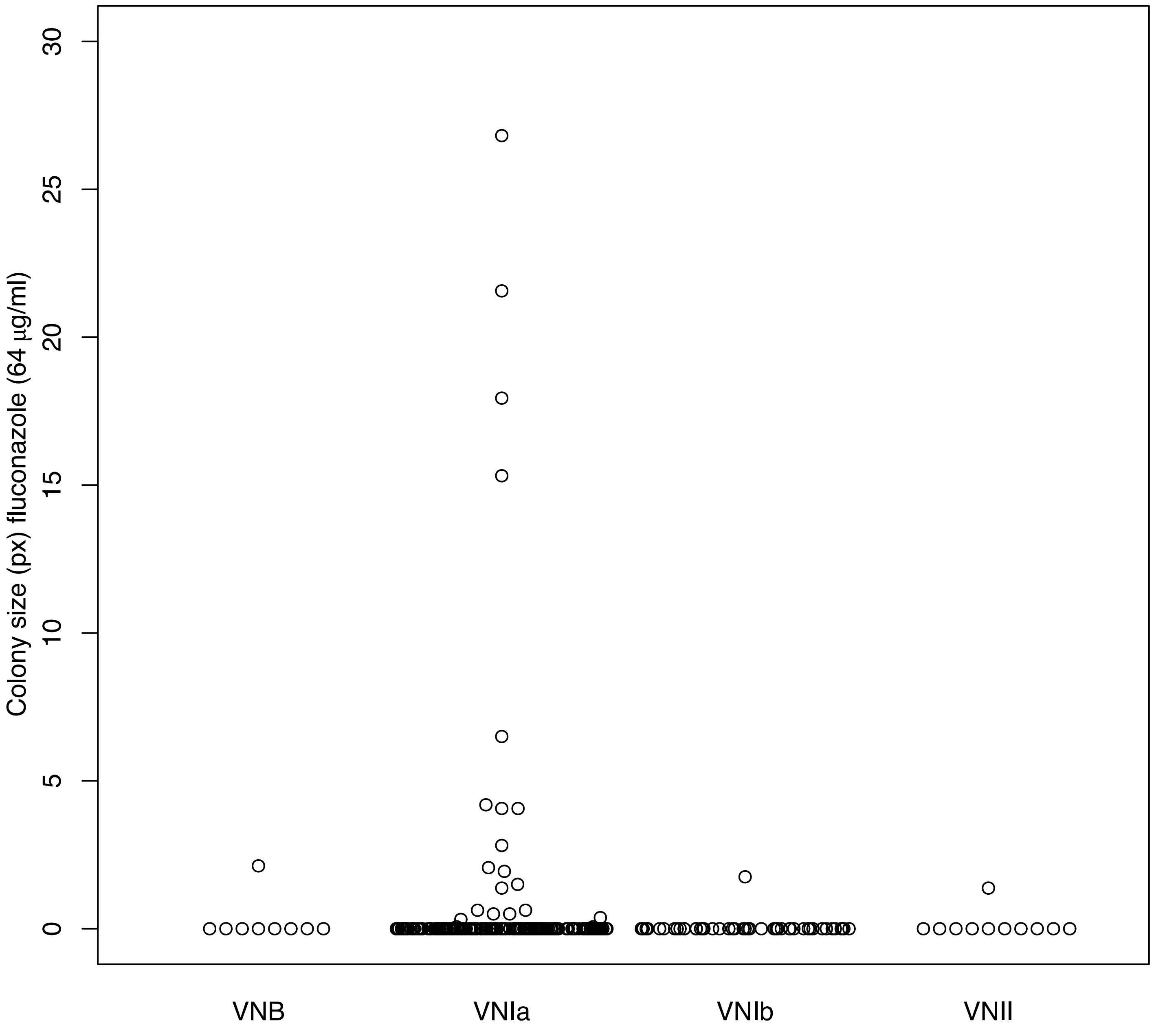


Supplemental Figure 3. Growth on fluconazole by lineage. Colony size (area, px) of isolates grown on fluconazole 64ug/ml for 3 days.
