## Supplemental Figure 4 for "Genomic variation across a clinical *Cryptococcus* population linked to disease outcome"

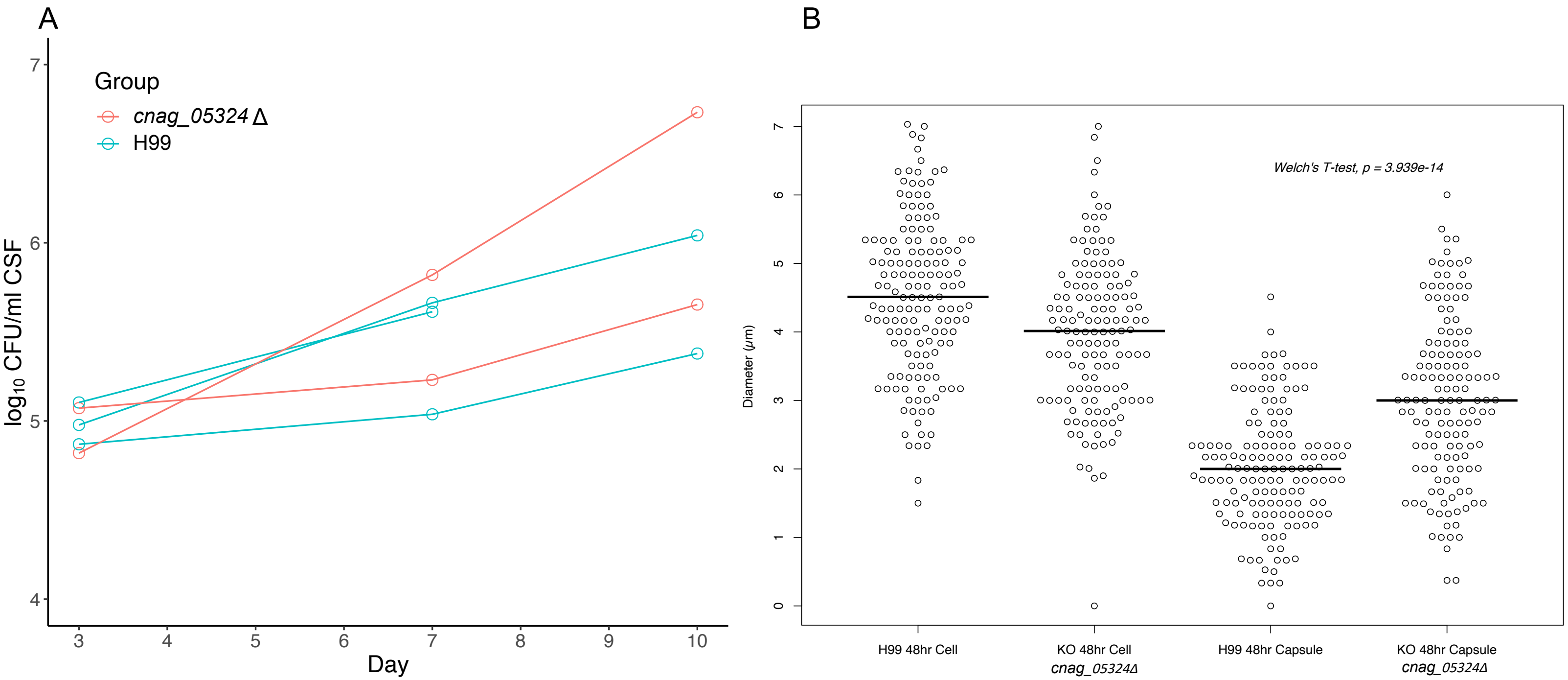


Supplemental Figure 4. Capsule and virulence regulation of CNAG_05324. A) Rabbit CSF load, Log10(CFU/ml), on days 3, 7, and 10, per rabbit (individual lines) infected with either H99 (blue) or the CNAG_05324 deletion strain (pink). B) Cell size and capsule diameter at 48 hours growth in capsule inducing media for H99 and the CNAG_05324 deletion strain.
