## Supplemental Figure 5 for "Genomic variation across a clinical *Cryptococcus* population linked to disease outcome"

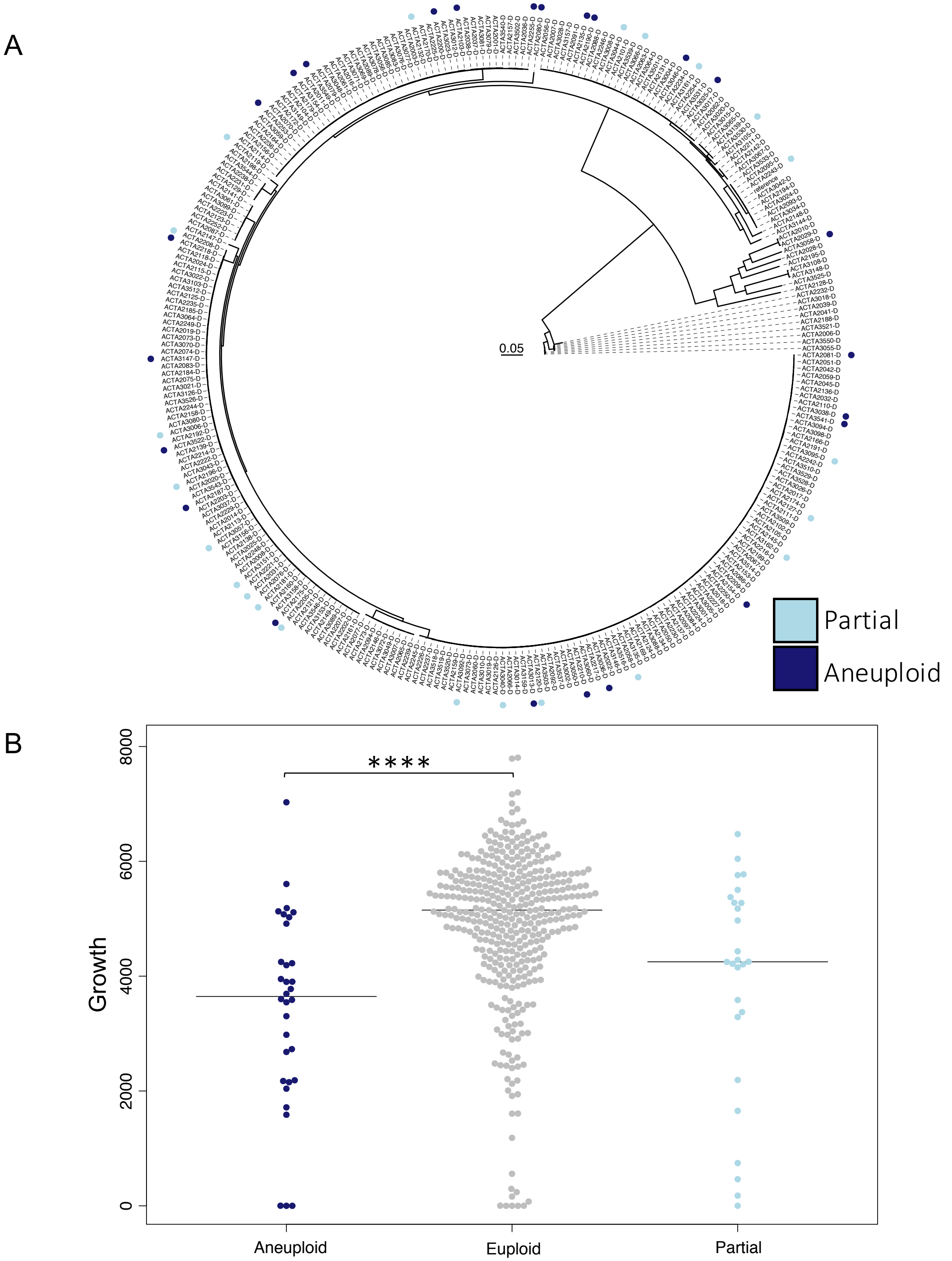


Supplemental Figure 5. Aneuploidy impact on growth for ACTA and Desjardins et al isolates. A) Whole (aneuploid) and partial chromosomal duplications throughout the population. B) Colony size (growth) by ploidy state on YPD at 37°C for both ACTA and Desjardins et al isolates.
