## Supplemental Figure 6 for "Genomic variation across a clinical *Cryptococcus* population linked to disease outcome"

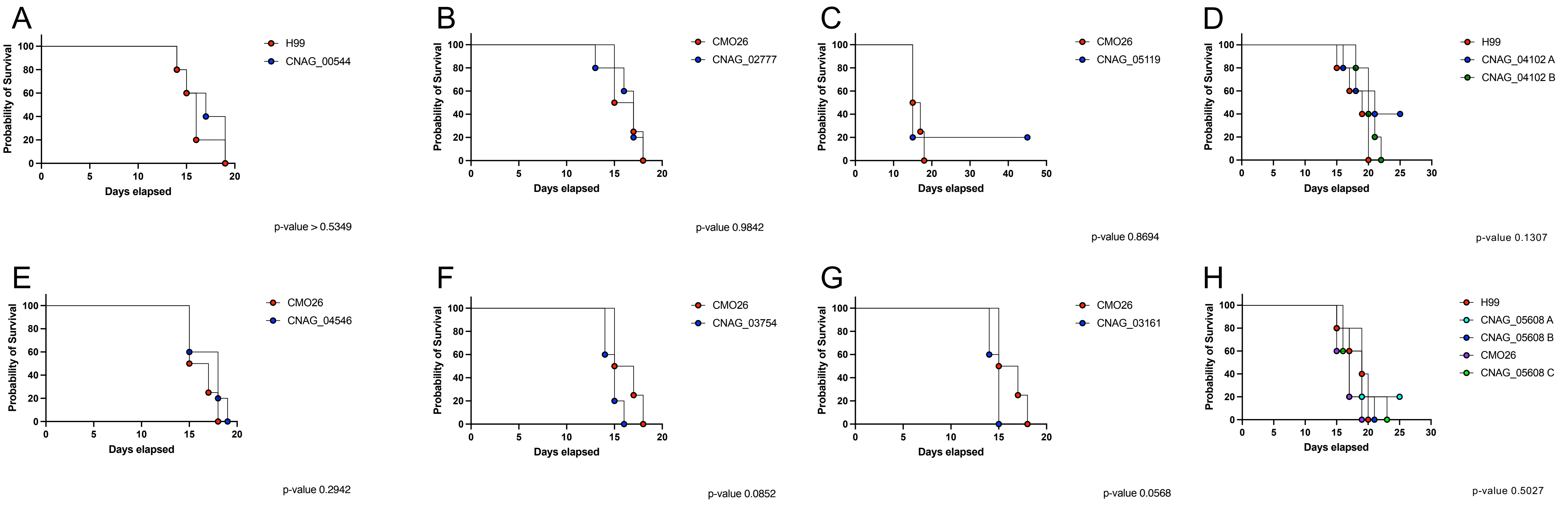


Supplemental Figure 6. Survival of mice infected with parental strain (H99/CMO26) or deletion strain. Five CD-1 mice were infected with approximately 5x10$^{4}$ CFU by oropharyngeal aspiration per strain. A) Survival of mice infected with parental strain (H99) and a CNAG_00544 mutant strain. B) Survival of mice infected with parental strain (CMO26) and a CNAG_02777 mutant strain. C) Survival of mice infected with parental strain (CMO26) and a CNAG_05119 mutant strain. D) Survival of mice infected with parental strain (H99) and two independent CNAG_04102 mutant strains. E) Survival of mice infected with parental strain (CMO26) and a CNAG_04548 mutant strain. F) Survival of mice infected with parental strain (CMO26) and a CNAG_03754 mutant strain. G) Survival of mice infected with parental strain (CMO26) and a CNAG_03161 mutant strain. H) Survival of mice infected with parental strain (H99/CMO26) and three independent CNAG_05608 mutant strains in the CMO26 background (C) and the H99 background (A, B).
